# Mechanisms underlying vaccination protocols that may optimally elicit broadly neutralizing antibodies against highly mutable pathogens

**DOI:** 10.1101/2020.10.07.330340

**Authors:** Raman S. Ganti, Arup K. Chakraborty

## Abstract

Effective prophylactic vaccines usually induce the immune system to generate potent antibodies that can bind to an antigen and thus prevent it from infecting host cells. B cells produce antibodies by a Darwinian evolutionary process called affinity maturation (AM). During AM, the B cell population evolves in response to the antigen to produce antibodies that bind specifically and strongly to the antigen. Highly mutable pathogens pose a major challenge to the development of effective vaccines because antibodies that are effective against one strain of the virus may not protect against a mutant strain. Antibodies that can protect against diverse strains of a mutable pathogen have high “breadth” and are called broadly neutralizing antibodies (bnAbs). In spite of extensive studies, an effective vaccination strategy that can generate bnAbs in humans does not exist for any highly mutable pathogen. Here we study a minimal model to explore the mechanisms underlying how the selection forces imposed by antigens can be optimally chosen to guide AM to maximize the evolution of bnAbs. For logistical reasons, only a finite number of antigens can be administered in a finite number of vaccinations; that is, guiding the non-equilibrium dynamics of AM to produce bnAbs must be accomplished non-adiabatically. The time-varying Kullback-Leibler divergence (KLD) between the existing B cell population distribution and the fitness landscape imposed by antigens is a quantitative metric of the thermodynamic force acting on B cells. If this force is too small, adaptation is minimal. If the force is too large, contrary to expectations, adaptation is not faster; rather, the B cell population is extinguished for reasons that we describe. We define the conditions necessary for the force to be set optimally such that the flux of B cells from low to high breadth states is maximized. Even in this case we show why the dynamics of AM prevent perfect adaptation. If two shots of vaccination are allowed, the optimal protocol is characterized by a relatively low optimal KLD during the first shot that appropriately increases the diversity of the B cell population so that the surviving B cells have a high chance of evolving into bnAbs upon subsequently increasing the KLD during the second shot. Phylogenetic tree analysis further reveals the evolutionary pathways that lead to bnAbs. The connections between the mechanisms revealed by our analyses and recent simulation studies of bnAb evolution, the problem of generalist versus specialist evolution, and learning theory are discussed.

## I. INTRODUCTION

Vaccines are a major contributor to the dramatic decline in childhood mortality over the past century, and have mitigated the threat of infectious diseases around the world. Indeed, vaccination has saved more lives than any other medical procedure. The importance of vaccines for the economy and for human health has been made vivid during the COVID-19 pandemic. Vaccines elicit pathogen-specific immune responses that can be rapidly recalled upon natural infection with the same pathogen, thus preventing disease. Most such prophylactic vaccines induce our immune systems to produce antibodies that can thwart infection by specific pathogens.

B lymphocytes (B cells), an important part of the immune systems of vertebrates, express a receptor on their surface that is called the B cell receptor (BCR). Humans have about 100 billion B cells each and most possess a BCR that is distinct from that of a different B cell. Antibodies are produced by a Darwinian evolutionary process called affinity maturation (AM) [1]. Upon stimulation by a pathogen or surrogate of a pathogen (antigen) that is used as a vaccine component, BCR on B cells can bind to proteins on the surface of the pathogen or antigen. B cells that bind with a free energy above a threshold can seed structures called Germinal Centers (GCs) where they undergo AM [2]. The BCR of these cells mutate at a high rate [3]. The mutated B cells compete with each other to bind to the pathogenic surface protein, and those with BCRs that can bind more strongly have a better chance to be positively selected. B cells that are not positively selected die. A few positively selected B cells exit the GC, and some of them secrete their BCR in soluble form. The secreted product is an antibody. Other positively selected B cells become so called memory B cells that can subsequently be stimulated rapidly upon exposure to the same pathogen. Most positively selected B cells undergo further rounds of mutation and selection. Therefore, as time ensues, the antibodies that are produced bind more strongly to the pathogenic surface protein [4].

To see how antibodies can prevent pathogens from infecting new cells, consider viruses. Viruses have spikes on their surface comprised of proteins. To infect a cell, a virus’ spike binds to a protein on the surface of the host cell. Antibodies can bind to the spike proteins of a virus and thus interfere with a virus’ ability to bind to a host cell protein, an action which prevents infection [5]. As a consequence, eliciting potent antibodies that bind specifically to any surface protein on the pathogen can be effective. Highly mutable pathogens can, however, present a major challenge to vaccination using this strategy. This is because mutations can emerge in their surface proteins, and the antibodies generated by vaccination, which bind to a particular protein, are no longer effective [6].

The surface proteins of even highly mutable pathogens, like HIV and influenza, have some regions that cannot mutate because these regions are important for binding to host cell proteins to enable infection [7–9]. For example, a part of the surface spike protein of HIV is relatively conserved. Antibodies that can bind to these conserved regions would be effective against diverse strains of the pathogens [10]. However, generating such antibodies by vaccination is a non-trivial challenge [11]. The relatively conserved region on the HIV spike is smaller in size than the typical size of an antibody’s antigen binding region, and the conserved part is surrounded by highly variable residues of the spike protein [12]. So, the challenge is to design HIV vaccination protocols that can guide the evolutionary process of AM to produce antibodies that bind only to the conserved bits of the spike and avoid the highly variable parts [13]. Antibodies that can achieve such specificity for the conserved parts of surface proteins, and can thus effectively neutralize diverse mutant virus strains, are called broadly neutralizing antibodies (bnAbs).

Antibodies that can bind to particular strains of a viral surface protein are specialists, while bnAbs may be viewed as generalists [14, 15]. AM is a stochastic nonequilibrium mutation and selection process. The challenge of influencing this process in a way that results in the evolution of generalists, rather than specialists, is a problem that lies at the intersection of non-equilibrium statistical physics, evolutionary biology, and immunology. Fundamental insights into this problem will be of significant pragmatic relevance to society.

Recently, simulation studies with the aim of shedding light on these issues have been carried out [2, 16–20]. A number of insights emerged from such studies. It is evident that in order to generate bnAbs, one would have to stimulate AM with multiple variant antigens that share conserved residues, but have different variable parts. Wang et al. [2] showed that sequential stimulation of AM with variant antigens, is likely a more effective way to generate bnAbs than a cocktail of the same antigens. Subsequently, experiments in mice have shown that antibodies resembling HIV bnAbs emerge upon sequential immunization with variant antigens [21, 22]. Upon vaccinating with one variant antigen, AM produces memory B cells that can be stimulated and undergo further evolution upon vaccinating with a second variant antigen, and so on. Each time a new variant antigen is introduced, the environment in which the existing memory B cells evolved changes, and the memory B cell population is driven from one steady-state to another [23]. So, sequential immunization results in the evolution of B cells by AM in a time-varying environment.

The non-equilibrium response of heterogeneous populations to time varying environments has been explored by numerous authors [24–26]. Clonal populations stochastically switch phenotypes in response to changes in the environment. Recently, stochastic phenotype switching was directed towards searching for optimal conditions that increase the fraction of generalists within a background population of both specialists and generalists [14, 15]. Two recent studies have provided further insights into how vaccination protocols could optimize the evolution of bnAbs, or generalists. Sprenger et al [17] have carried out simulations of the AM process upon sequential immunization and reported that there is an optimal distance between the variable region of the immunogens used in sequence, and/or immunogen concentration, in order to maximize the chance of producing bnAbs. Furthermore, they found that the optimum distance or concentration is sequentially higher as each new variant antigen is used to stimulate AM. The optima were thought to correspond to optimal extents to which the existing B cell population is driven out of equilibrium from a previous steady state. Recently, Sachdeva et al. [15] studied the effect of cycling between antigenic environments. Their simulations showed that if a population of specialist B cells is subjected to time-varying environments cycled at an optimal resonance frequency, specialists will evolve to become generalists while the reverse process is prevented. Interestingly, it was observed that gradually increasing the frequency from slow to fast resulted in an optimal point for generating a large fraction of generalists. This result is related to the shift of the optimum to higher values at each subsequent step of sequential immunization that Sprenger et al. reported [17].

The goal of vaccination is to maximize the production of bnAbs. One of the practical constraints is that we are allowed only a small finite number of immunizations due to logistical considerations. From a theoretical standpoint, this implies that the selection forces cannot be applied in an adiabatic fashion to guide the evolutionary dynamics to the desired end point. The non-adiabatic character of the imposed selection forces with the possibility of B cell death (absorbing boundary condition) represents an interesting problem in non-equilibrium statistical mechanics.

Previous studies of this problem have considered the effects of changing immunization protocols by altering the mutational distance and concentrations of antigens [2, 16, 17, 27] or varying the frequency of environmental cycling [15]. Here, we have formulated the problem so that the effects of changing these variables can be clearly related to concepts in non-equilibrium statistical mechanics and learning theory. We provide a quantitative measure of the effective thermodynamic selection force imposed on the B cell population upon immunization, and explore how varying this force influences when and how the B cell population adapts to the selection force. We use this conceptual framework to provide insights into the evolutionary forces that define optimal vaccination protocols. The key features of evolutionary trajectories that lead to the evolution of generalist bnAbs are also described.

## II. RESULTS

### A. Minimal model for the evolution of broadly neutralizing antibodies

As we described in our introductory remarks, upon natural infection or vaccination, antibodies that bind more strongly to the stimulating antigen evolve by a Darwinian evolutionary process. Our focus here is the evolution of bnAbs, which bind to regions of the virus’ surface proteins that are relatively conserved across strains.

The virus’ intact surface protein complex (e.g., the spike), or a part of it, has to be the immunogen, or the active component of the vaccine. This is because the B cells that evolve during AM must learn to focus their binding footprint on the relatively conserved residues in the context of the actual geometry of the surface protein. But, the protein has many regions that do not contain any conserved residues. Human B cells have a huge diversity of BCRs, and so vaccinating with the surface protein would most likely result in activating B cells that do not bind to the region containing conserved residues [11]. This is because there are many more regions without conserved residues. AM would then proceed, and generate antibodies that bind well to the variable residues of a particular strain of the surface protein that was employed in the vaccine. To address this challenge, immunogens have been developed that can activate the so called germline B cells that bind to parts of the protein containing the conserved region [10]. These immunogens are not the surface protein complex, but something much simpler. Now, upon vaccination with mimics of the full surface protein complex, can bnAbs evolve? If so, what is the optimal strategy? These are the questions we consider here in terms of non-equilibrium thermodynamic forces and fluxes. Our findings can help guide practical choices regarding vaccination protocols in terms of concrete concepts in statistical physics and learning theory.

To obtain essential physical insights into this problem, we construct a minimal model. The affinity between a BCR and an antigen to which it binds is, in principle, defined by many variables, such as the character of the amino acid residues of the BCR’s antigen binding region and those of the viral epitope as well as the threedimensional conformations of the interacting parts. Various coarse-grained models have been used to represent such interactions. One class of such models described by Wang et al. [2], Luo et al. [27], and Sprenger et al. [17] represented the antigen binding region of the BCR and the epitope using strings of sites, and used simplified models to calculate binding free energies between them. The probability of a B cell being positively selected depended on this interaction free energy. Another simplified representation, called shape space, has proven useful for reducing the dimensionality down to a few (5 or 6) abstract variables [28–30]. Recently [16], a 2D model of shape space was used to simulate GC reactions, where one of the dimensions corresponded to conserved amino acids and the other to variable amino acids on the antigens. The BCR regions that interact with the conserved and variable amino acids of the antigen were represented similarly on the two axes. The affinity depended on the Euclidean distance between the BCRs and antigens on a hypersphere centered around the origin. Similarly, the breadth of evolving BCRs will be defined by many variables. Inspired by the shape-space model, and in the spirit of Occam’s razor, we consider the breadth of coverage of a BCR to be defined by a single dimension. This dimension may be considered to be an appropriate projection of a higher-dimensional manifold, and we will refer to it as the “breadth space”.

The origin (0.0 on the abscissa of (Fig. 1(a)) denotes the state of highest possible breadth. If a B cell with a particular BCR sequence is at the origin, it binds with the highest possible affinity to conserved epitopes on the antigens and avoids binding to the surrounding variable regions as best as possible. B cells traverse the breadth dimension by mutations and the breadth decreases upon moving left or right from the origin. We discretize breadth space into a set of *K−*1 bins where *K* is the total number of states and the additional state corresponds to death (Fig. 1(b)). Mutations that result in changes of affinity between BCR and antigens, and thus changes of the breadth state in our model, occur because of discrete modifications to codons that code for BCR amino acids. Also, changes in breadth state are the product of multiple mutations [31, 32]. Thus, we believe that a discrete representation of breadth space is appropriate. Mutation-induced change in affinity between proteins (i.e., BCR and antigens) are log-normally distributed [17]. We account for this by setting the mutational transition probabilities such that breadth-enhancing mutations are less likely to occur than breadth-reducing mutations. Some mutations can result in a BCR that does not fold into the proper shape or confer some other grossly deleterious feature. So, any BCR sequence, with some probability, can acquire a lethal mutation [33]. Also, GC B cells are intrinsically apoptotic; i.e., if not positively selected, they die [34, 35].

**FIG 1:**
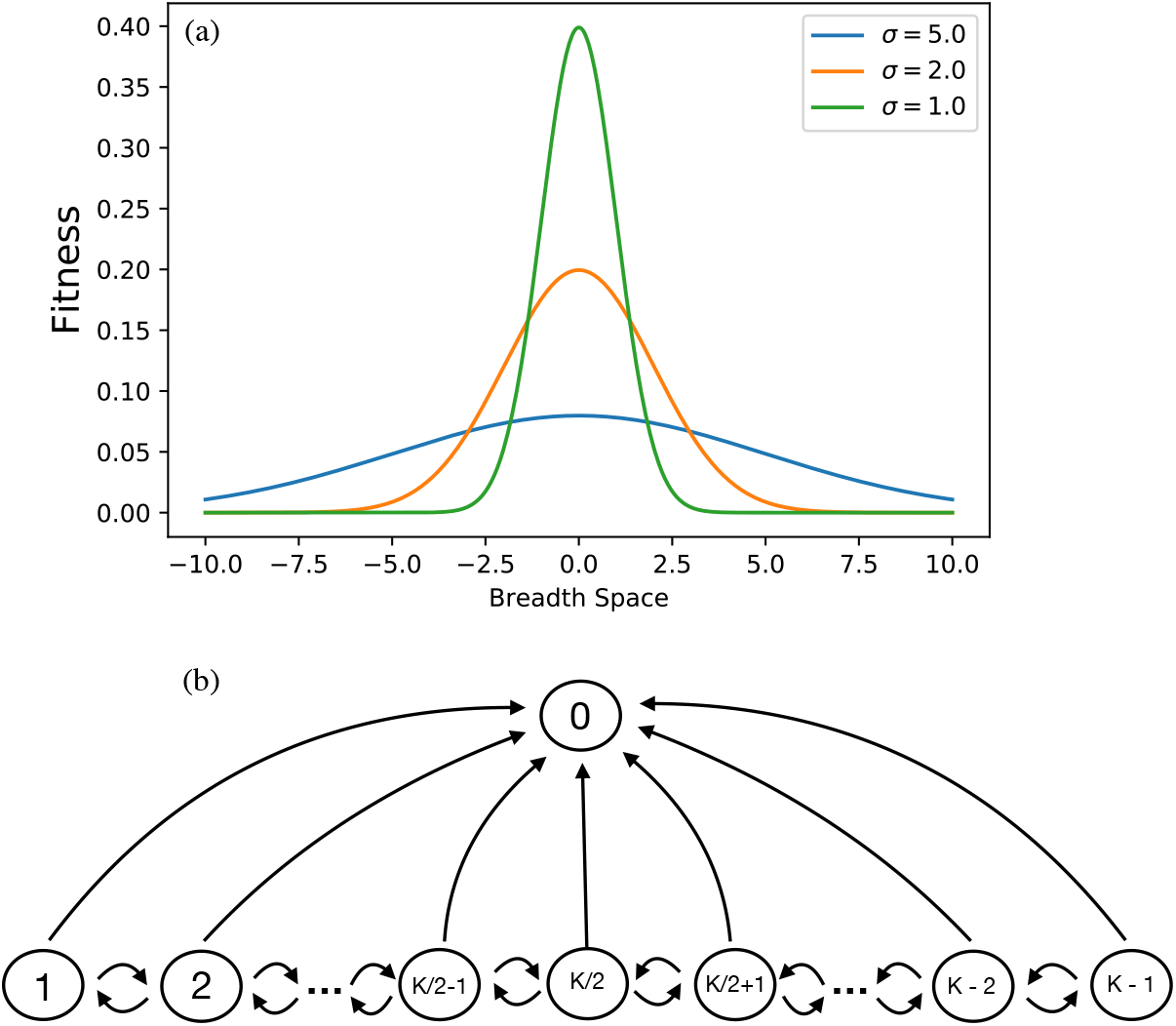
Definition of breadth space. (a) Depiction of the 1D fitness landscape. The origin denotes the point of highest breadth and the fitness distribution imposed on the B cell population by the immunogen is Gaussian. Reducing the variance of the fitness focuses selection pressure on B cells that bind with high affinity to conserved epitopes. (b) Breadth space is binned into *K −* 1 states: *K/*2 is the highest breadth state corresponding to the origin in (a), 1 and *K −* 1 are states of lowest breadth, and 0 is the death state.

The immunogen that leads to the activation and maturation of B cells that can bind to the conserved residues on the surface protein produces a pool of memory B cells that have the potential to evolve into bnAbs. However, these B cells are not bnAbs. The immunogens that mimic the entire virus’ surface protein and are administered subsequently are meant to evolve this B cell population to produce bnAbs. These immunogens, the variant antigens that share conserved residues, impose selection forces on this population of B cells.

Different B cells correspond to different states of “breadth”. The selection force imposed on a particular B cell depends upon its breadth. During affinity maturation, B cells compete with each other to be positively selected by T helper cells that are present in limiting numbers. The probability of a B cell in a particular breadth state (or bin) being positively selected per unit time can be regarded as the “fitness” of B cells in that bin. So, we will refer to this probability as fitness hereon.

Since the goal is to maximize the number of B cells occupying the highest breadth state, the selection probability or fitness landscape is taken to be a Gaussian centered at the bin corresponding to bnAbs, as shown in Fig. 1(a). The choice of immunogen and immunization protocol determines the variance of the imposed fitness landscape. For example, if a cocktail of variant antigens that share conserved residues but are very different in their variable regions, is administered, only the B cells that have evolved high breadth will be strongly selected. The corresponding fitness landscape will be sharply peaked (orange and green curves in Fig. 1(a)). If a single immunogen is first administered, then a greater diversity of B cells can be positively selected, and the fitness landscape is characterized by a higher variance (blue curve in Fig. 1(a)). For sequential immunization, as noted above, the fitness landscape changes in discrete time steps. This change corresponds to a time-varying antigenic environment in which a heterogeneous B cell population evolves through replication, mutation, and selection.

The birth-death master equations describing time evolution of the probability distribution of the B cell population vector subjected to mutation and selection is given by [25]:

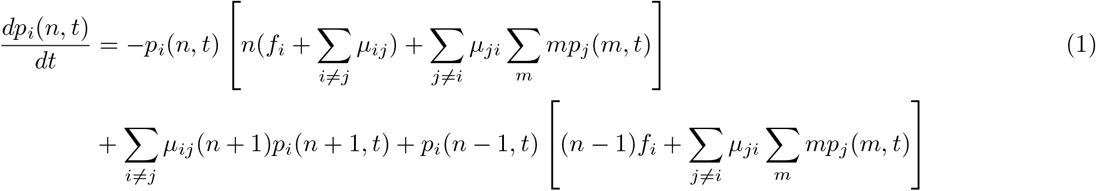

where *i* denotes the index of the breadth bin, *n* and *m* are the number of B cells in bins *i*, and *j*, respectively; *µ*_*ij*_ is the mutation rate per cell from bin *i* to *j, f*_*i*_ is the fitness or probability per unit time that a B cell in bin *i* replicates. The combined effects of lethal mutation and basal death rate of GC B cells corresponds to a rate of B cell death, denoted by *µ*_*i*0_. The state space describing the master equations is shown in Fig. 1(b). B cells that occupy bin *K/*2 have attained maximum possible breadth whereas those in states 1 and *K−* 1 have lowest breadth. State 0 represents an absorbing boundary condition as it corresponds to B cell death.

Directly solving the master equations is numerically cumbersome. Instead, taking the expectation value of Eq (1) and following the procedure described in the SI of [25], we derive the following mean-field equations:

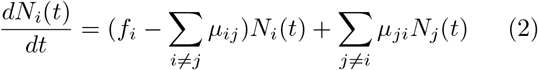

where *N*_*i*_ is the number of cells occupying bin *i* and *N*_*j*_ is the number in bin *j*. Eq (2) clarifies the processes which regulate the population dynamics in each breadth state. Cells in a particular bin replicate with a probability per unit time that depends on the fitness of cells in the bin and the number of cells that occupy that bin. They leave bin, *i*, with a propensity equivalent to the product of the transition rate from *i* to all other states *j* and the population of cells in bin, *i*; entrance into bin, *i*, is determined by rates *j* to *i* and occupancy of states *j*. Of course, if *i* = 0, the first term on the right-hand side of Eq (2) vanishes.

The master equations (Eq (1)) can be solved by transforming the mean field equations (Eq (2)) into a set of “chemical reactions” and using the Gillespie method [36, 37] (see Supplementary Materials) to generate stochastic trajectories of the B cell birth-death-mutation process; each trajectory contains the set of all reactions that occur within a single GC. There are two important stop conditions that end a GC trajectory: the B cell population dies (Σ _*i* ≠ 0_ *N*_*i*_(*t*) = 0) or the total number of B cells in the GC approaches a sufficiently large size. The former condition represents extinction, and the latter is a proxy for the B cells having consumed all antigens present in the GC, thus ending AM. The objective of our stochastic simulations is to understand how a discretized time-evolving fitness profile or changing antigenic environment affects the production of bnAbs, and thus determine the mechanistic underpinnings of how certain vaccination protocols may optimize bnAb production. In experimental studies [38], antigen concentration and mutational distance between the variable regions of sequentially administered immunogens are the principal parameters that can be controlled by the vaccine administrator. Changing these parameters changes the selection probability and therefore the characteristics of the fitness landscapes. In this case, a large decrease in the antigen concentration or large increase in the mutational distance corresponds to a large decrease in the variance of the fitness landscape.

In response to an immunogen, the existing memory B cell population evolves with respect to the imposed fitness landscape (Fig. 2(b), orange). The amount of information that the B cell population needs to gain in order to adapt perfectly to the antigenic environment is quantified by the Kullback-Leibler divergence (KLD) or the relative entropy of the B cell population distribution after injection *j* (*p*^*j*^) and the fitness imposed (probability of selection per unit time) during injection *j* + 1 (*f*^*j*+1^) [39]:.

**FIG 2:**
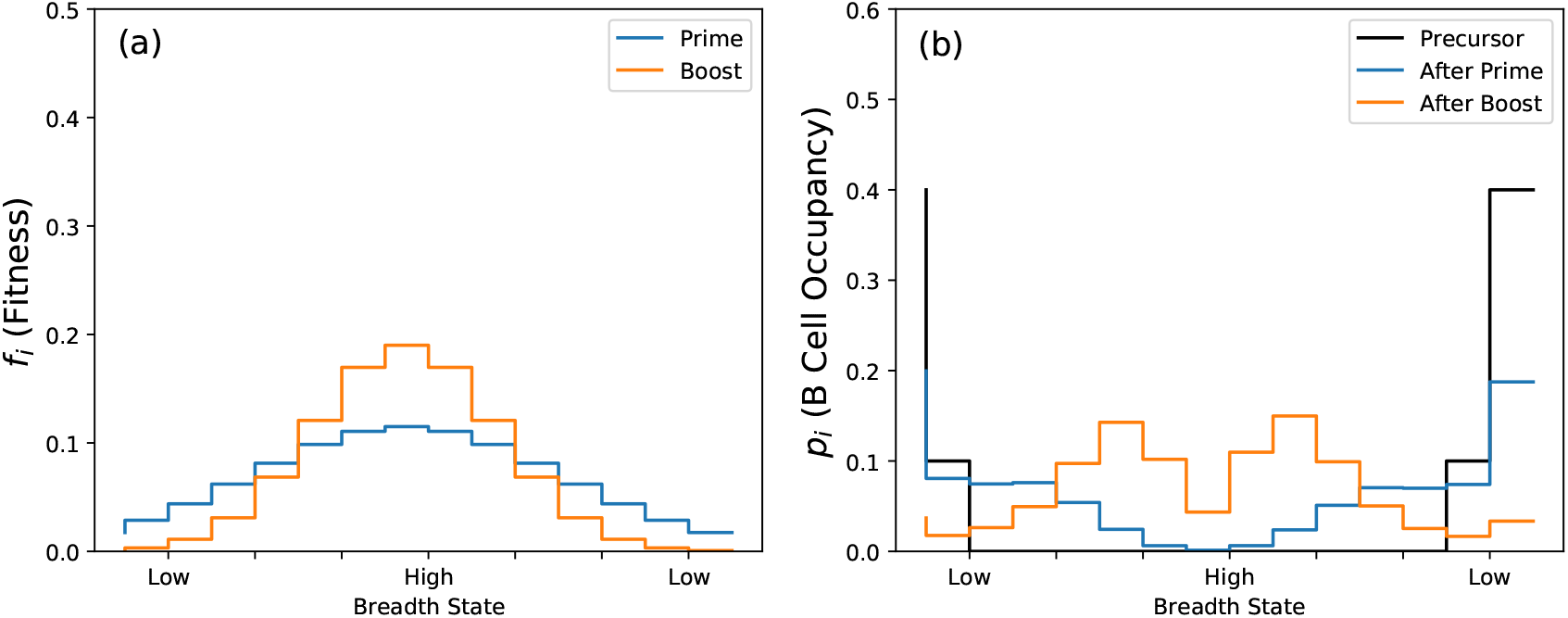
Fitness landscape and response of the B cell population. (a) Imposed fitness landscape (*f*_*i*_) during prime (blue) and boost (orange). (b) Response of B cell population distribution *p*_*i*_ to fitness landscapes shown in (a).

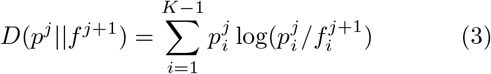

Eq (3) quantifies the thermodynamic force acting on the existing B cell population upon immunization: the KLD between the probability distribution describing the breadth states of the B cell population and the distribution describing the probability with which states of different breadth are positively selected per unit time (i.e., fitness). In the context of statistical learning theory, Eq (3) defines a cost function that must be minimized by an evolving B cell population. Note also that the KLD as defined in Eq (3) is zero when the steady state corresponding to extinction (*p→*0) is approached. The quantity *D*(*f*∥*p*) would approach infinity in the same limit, which is undesirable. A symmetrized form of the KLD e.g. the Jensen-Shannon divergence would have a similar undesired behavior. To study the effects of varying the immunization protocol, we will vary the KLD.

The parameters, *µ*_*ij*_ and *µ*_*i*0_, must be fixed so that the resulting dynamics are consistent with the expected behavior of GCs and B cell sequence evolution [17]. Given that breadth space is discretized into *K−*1 bins and the fitness landscape is normalized, *µ*_*i*0_ *<* 1*/*(*K−*1) so that the population does not become extinct when a uniform fitness profile is imposed. The fitness landscape, not the basal death rate, should primarily determine the probability of GC survival. It is known from clinical data on HIV that the initially activated B cells need to acquire many mutations in order to become bnAbs [31, 32]. Thus, even when bnAbs evolve naturally, it usually takes a long time. In our coarse-grained minimal model, this observation translates to a low mutation rate between bins. This is because each mutation in our model corresponds to many mutations at the residue level. Also, breadth-enhancing mutations are far less likely to occur than those that reduce breadth [40]. In biological terms, this is because B cells are more likely to mutate away from the small number of sequences that have high affinity for the conserved residues than to mutate towards them. In other words, the sequence entropy of high-breadth B cells is lower than that of low-breadth cells; as a consequence, an entropic force pushes B cells to mutate outward from low (high breadth) to high (low breadth) sequence entropy states.

An initial population of 50 cells is sampled from the occupancy distribution (black, Fig. 2(b)). These are the B cells that were activated by the simple immunogen that selects for cells that bind to the region of the virus’ protein that includes the conserved residues [38, 41]. As noted earlier, the selected B cells are not bnAbs, which is why the initial occupancy distribution of B cells is chosen as shown in Fig. 2(b). A fitness landscape is then imposed on this B cell population (blue, Fig. 2(a)), which corresponds to a particular KLD (Eq. (3)). As a consequence, B cell occupancy for all GCs that did not go extinct shifts from low to higher breadth bins (blue, Fig. 2(b)). We run 100 GC simulations, and obtain stochastic trajectories of the reactions derived from Eq (2) until either stop condition is met: *N*_*total*_ = 0 or 200. From each surviving GC, 50 new memory B cells are sampled from the 200 that exit after prime. Corresponding to the second immunogen, a new fitness profile (KLD) is imposed and this B cell population then evolves stochastically until either stop condition is met. The procedure can be repeated for subsequent immunizations, and the total number of B cells occupying the highest breadth state (*i* = *K/*2) summed over all surviving GCs divided by the number of GCs can be regarded as a measure of the average number of bnAbs produced by the immunization protocol. For the results that are shown, *K* = 16, *µ*_*i*0_ = 0.02, and *µ*_*ij*_ = 0.05. The mutation rates *µ*_*i,i*+1_ = 0.125*µ*_*ij*_ and *µ*_*i*+1,*i*_ = 0.875*µ*_*ij*_ if *i < K/*2. Conditions flip for *i > K/*2 and for *i* = *K/*2, *µ*_*i,i*+1_ = *µ*_*i,i−*1_ = 0.5*µ*_*ij*_.

### B. Optimal protocols for the first and second immunizations

We will focus here on studying vaccination protocols with two sequentially administered immunogens. Following standard terminology from vaccination, we will refer to the first as “prime”, and to the second as “boost”. There is a large continuous space of choices for immunogens, or possible fitness landscapes, for prime and boost. We first asked if there is an optimal combination of the imposed fitness landscapes, *f* ^1^ and *f* ^2^, respectively, that maximizes bnAb production. Since the fitness landscape corresponding to the boost immunogen acts on the B cell distribution that evolved in response to the fitness profile corresponding to the priming immunogen, it is clear that optimal *f* ^2^ will depend on the choice of *f* ^1^. Using the previously described parameter values, we first searched over the space of variances in *f* ^1^ profiles that are imposed on the B cell population (*p*^0^) that developed after activation of the right germline B cells. We then searched over the space of variances of *f* ^2^ profiles that can be imposed on *p*^1^, the distribution that evolves after prime immunization. Our results (Fig. 3(a)) show that an optimal setting of *f* ^1^ exists (yellow line), occurring near a KL divergence *D*(*p*^0^∥*f* ^1^) = 2.76.

**FIG 3:**
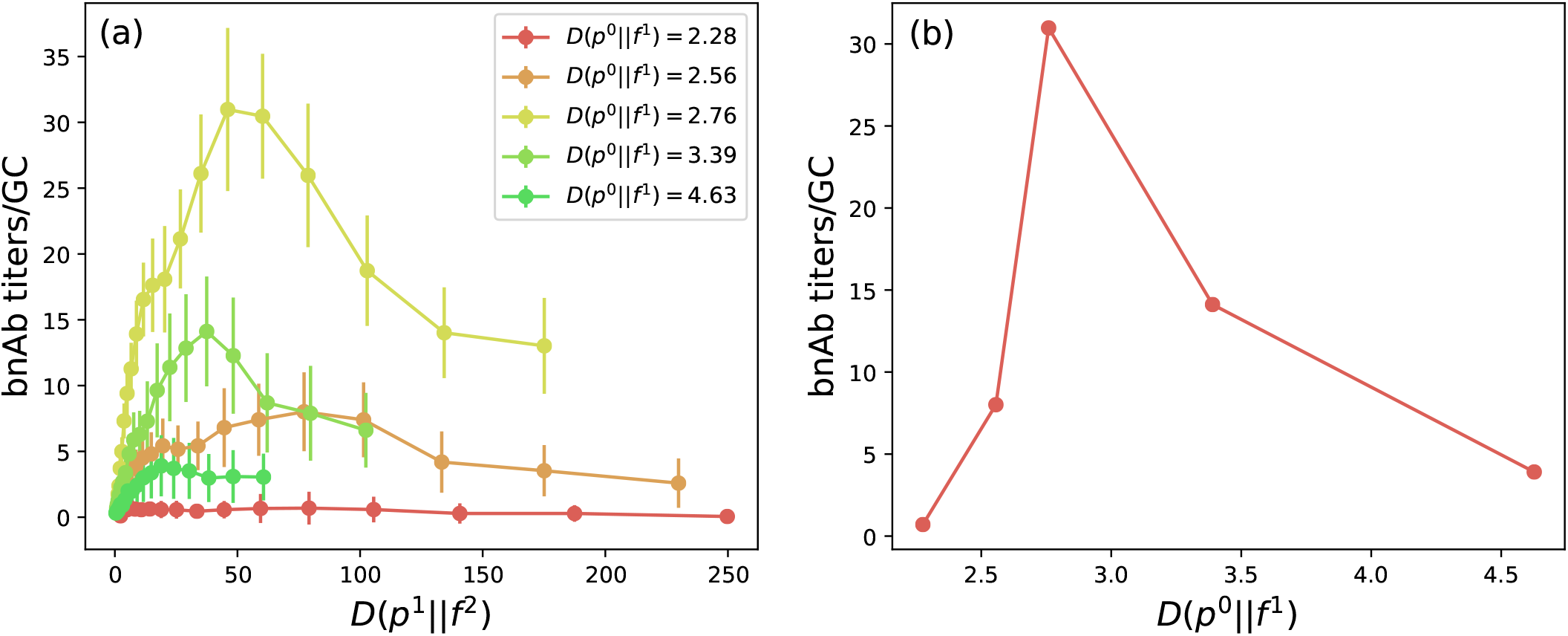
Response to different vaccination protocols. (a) For each prime immunization choice of KL divergence, *D*(*p*^0^∥*f* ^1^) (colored curves), the number of bnAbs produced per GC is shown over a range of boost immunization choices, *D*(*p*^1^∥*f* ^2^). (b) Maximal bnAb titers for each of the curves shown in (a) versus *D*(*p*^0^∥*f* ^1^) shows that an optimal prime-boost protocol exists.

For each choice of *f* ^1^, the number of bnAbs produced per GC after boost is plotted as a function of *D*(*p*^1^∥*f* ^2^), the divergence between the boost fitness *f* ^2^ and the B cell distribution that results after prime *p*^1^ (Fig. 3(a)). In all cases, there is also an optimal setting of *D*(*p*^1^∥*f* ^2^), which exceeds *D*(*p*^0^∥*f* ^1^). From the standpoint of vaccination, the last result implies that boost immunization needs to more aggressively focus selection on high breadth B cells than prime in order to maximize production of bnAbs. This result is consistent with what has been found with more elaborate computer simulations that reported an optimal mutational distance between immunogens and antigen concentration or oscillatory frequency of environmental changes [15, 17].

At these optimal points, we computed the probability of GC survival; i.e., the probability that the B cell population is not extinguished during AM. Fig. 4(a) shows that P(GC Survival) = 1.0 for low values of *D*(*p*^0^∥*f* ^1^) during prime immunization. At the optimal point (*D*(*p*^0^∥*f* ^1^) = 2.76), P(GC Survival) drops to*∼* 0.9. Beyond the optimal value of *f* ^1^, there is a sharp drop in GC survival probability (Fig. 4(a)) and bnAb titers (Fig. 3(b)). Fig. 4(b) provides an explanation for the abrupt onset of GC death past the optimal point. For low KL divergences (*D*(*p*^0^∥*f* ^1^) *≤* 2.56), the fitness of all B cell breadth states exceeds the intrinsic death rate *µ*_*i*0_ = 0.02 (blue dotted line, Fig. 4(b)). As a result, B cells sampled from the precursor population (black, Fig. 2(b)) that predominantly occupy low breadth states proliferate quickly, leading to GC survival. As these B cells rapidly internalize antigen, *N* quickly reaches a value of 200, and AM ends. Thus, there is limited time for mutations that allow B cells to transition from low to high breadth states.

**FIG 4:**
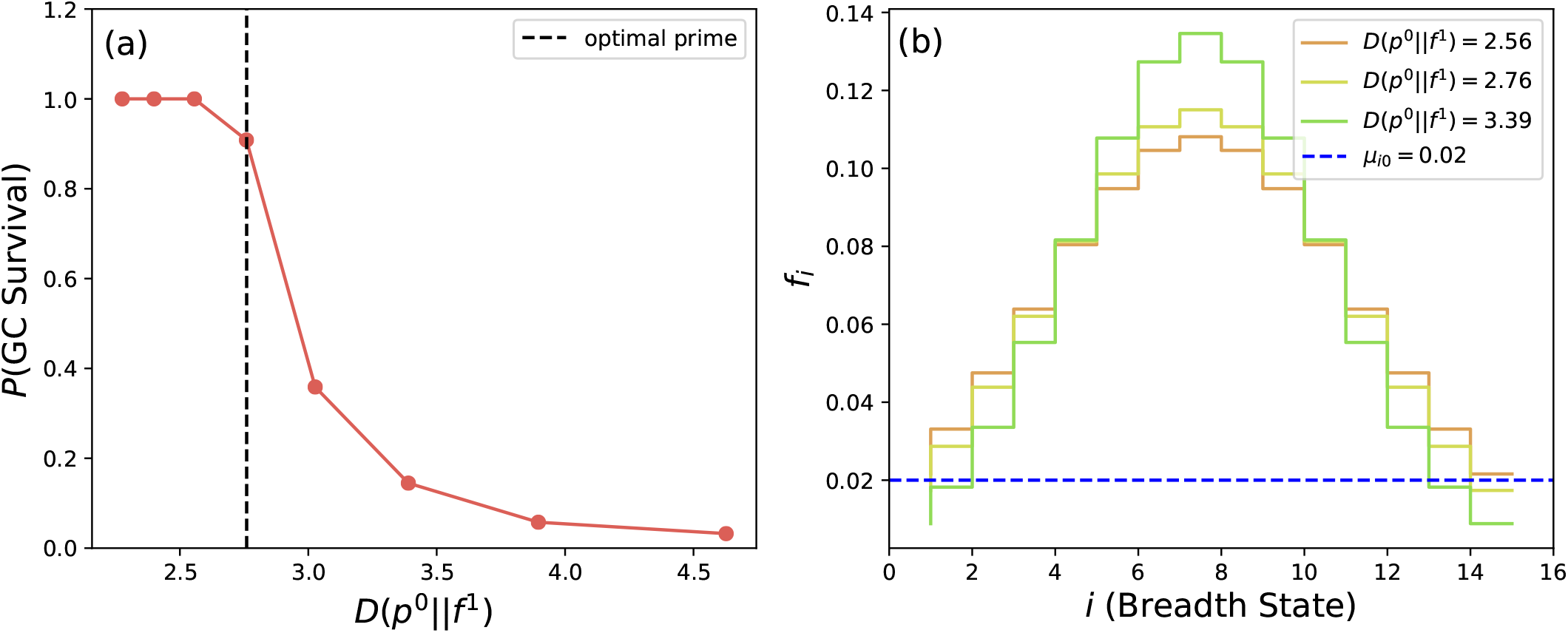
Germinal center (GC) survival (B cell population is not extinguished). (a) Fraction of GCs that survive prime immunization for a range of values of *D*(*p*^0^∥*f* ^1^). The fraction falls to*∼* 0.9 at the optimal setting and drops significantly beyond this point. (b) Sample fitness landscapes at the points located on the curve shown in (a).

For *D*(*p*^0^∥*f* ^1^) *>* 2.76, we observe that fitness in the lowest breadth states drops below the death rate. Furthermore, the occupancy of B cells in states adjacent to the bnAb state is low. So, very few of these B cells multiply, and most mutations that arise during proliferation are transitions to lower breadth states. Mutational flux to low breadth leads to accumulation of B cells in states at the edge where they die. As a consequence, there is a sharp increase in the likelihood of extinction events when the fitness of the lowest breadth states drops below the death rate. At *D*(*p*^0^∥*f* ^1^) = 2.76, the fitness of the lowest breadth states (yellow, Fig. 4(b)) drops just below the death rate. But, the B cells in states further from the edge states have reasonable fitness and can replicate. Some of these cells mutate toward higher breadth states and replicate more, while others transition to low breadth states and die. This leads to an optimal balance of these fluxes that causes neither rapid extinction nor proliferation that could end the AM process. Under these conditions, GC reactions continue for some time, enabling the B cell population to acquire the many mutations required to become bnAbs.

In order to rule out any effects due to coarse-graining of breadth space, additional calculations with a larger number of bins were carried out and the results are shown in the Supplementary Material. For *K* = 32 and using the same assumptions regarding the parameter regime, we observe the same qualitative behavior (Figs. S1-3).

### C. Understanding the optimal conditions in terms of thermodynamic forces and fluxes, and learning theory

Previous studies have used the KLD to quantify the distance between equilibrium distributions of phenotypic traits [42]. Upon a new immunization (prime or boost), if the KLD at this time is not too large, the B cell population evolves in an attempt to adapt to the imposed selection force, resulting in reduction of the KLD with time. Thus, as Fig. 5 shows, after each injection, the time-dependent B cell population distribution evolves to reduce the KLD, indicating that B cells learn about the environment through mutation and selection. However, perfect adaptation to the imposed selection forces (*D*(*p*∥*f*) = 0) does not occur; when the B cell population reaches the maximal threshold and AM ends, the KLD equals a finite positive value (Fig. 5). Note that *D*(*p*∥*f*) = 0 also corresponds to an equilibrium state. Taken together, these findings show that, if the thermodynamic force driving adaptation is not too large, the B cell population reaches a non-equilibrium steady state. Perfect adaptation is possible if the population continues to expand. Fig. S4 shows that if the population were allowed to increase indefinitely, corresponding to continuous antigen stimulation, the system eventually relaxes to a near equilibrium state (*D*(*p*∥*f*) *∼* 0). This is the adiabatic limit, which is unrealistic for discrete immunizations. It is noteworthy, however, that experiments and analyses have shown that slow delivery of immunogens over a time period improves affinity maturation [43, 44].

**FIG 5:**
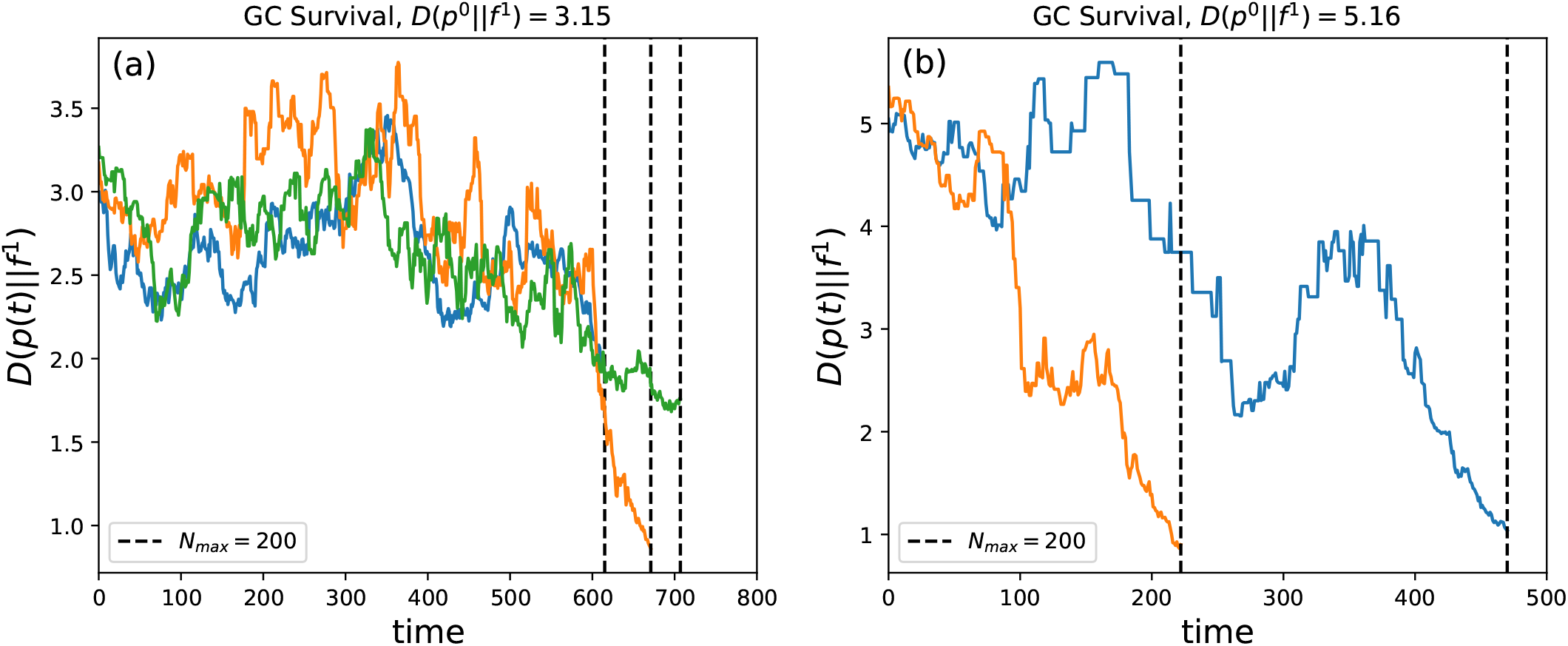
Learning inside a GC. Within surviving GC trajectories, the system eventually approaches a regime where *D*(*p*(*t*)∥*f* ^1^) decreases. On average, the rate of decrease is slower (a) near optimal forcing (*D*(*p*^0^∥*f* ^1^) = 3.15) and faster (b) beyond the optimum, *D*(*p*^0^∥*f* ^1^) = 5.16.

Typically, non-equilibrium thermodynamics tells us that if the thermodynamic force is larger, adaptation should occur faster. Indeed, this does happen in rare evolutionary trajectories when KLD is very large (Fig. 5(b)). However, if the initially imposed KLD upon the prime or boost immunization is too large, the B cell population is extinguished with high probability. This is because very few, if any, B cells in the population have a significant probability of being positively selected shortly after the new selection force is imposed and there is a net mutational flux to states of lower breadth (and probability of positive selection). Thus, the dynamical system acquires a steady state corresponding to the absorbing boundary condition of B cell death and extinction.

In the context of statistical learning theory, reduction of the KL divergence (Eq (3)), or the cost function, indicates that learning is happening. Fig. 5 shows that in GC trajectories where the population survives, *D*(*p*(*t*) ∥*f* ^1^) decreases over time until the maximum population size (*N*_*max*_ = 200) is reached. If a selection force near the optimal point is exerted on the B cell population (Fig. 5(a)), the KLD decreases more slowly than if a strong selection force is imposed (Fig. 5(b)). The KLD only decreases if breadth-enhancing mutations allow B cells to transition to higher breadth states. Under strong selection forces, if B cells do not mutate fast enough to higher breadth states, they rapidly die and extinction occurs. The few GC trajectories that survive (shown in Fig. 5(b)) do so because breadth-enhancement and therefore learning happens quickly.

In a conventional supervised learning problem, the cost function is initially set by the distance between the data and model distributions [45]. The learning rate during gradient descent must be chosen optimally. If it is too low, training will happen slowly and the model distribution will take a long time to fit to the data distribution. If it is set too high, undesirable divergent behavior can emerge and the cost function can increase over time. As the variance of the fitness distribution changes upon choosing different immunogens, the learning rate per injection is set by *D*(*p*^*j*^∥*f*^*j*+1^). If it is too low, the population increases rapidly before any substantial decrease in KLD is observed. If it is too high, most of the GC trajectories die. In the trajectories that lead to extinction (Fig. S5), we clearly observe divergent behavior as the KLD may increase rapidly before the population dies.

### D. Optimal KL divergence during the prime generates the right diversity of B cells for subsequent evolution of bnAbs

After prime immunization, as the B cell population acquires mutations that confer higher breadth, it diversifies to occupy bins corresponding to higher breadth states. Thus, *D*(*p*∥*f* ^1^) decreases. Fig. 6(a) shows a histogram of the relative entropy change (Δ*D*(*p*∥*f* ^1^)) of the B cell distribution within all GCs. Δ*D*(*p*∥*f* ^1^) is calculated by computing the difference between the KLD just before GC exit and the initially imposed value, *D*(*p*^0^∥*f* ^1^). At low *D*(*p*^0^∥*f* ^1^), all GCs survive but produce B cell populations dominated by low breadth cells (orange, Fig. 6(b)). As a result, in the majority of GCs, there is a minimal decrease in the KLD (orange, Fig. 6(a)).

**FIG 6:**
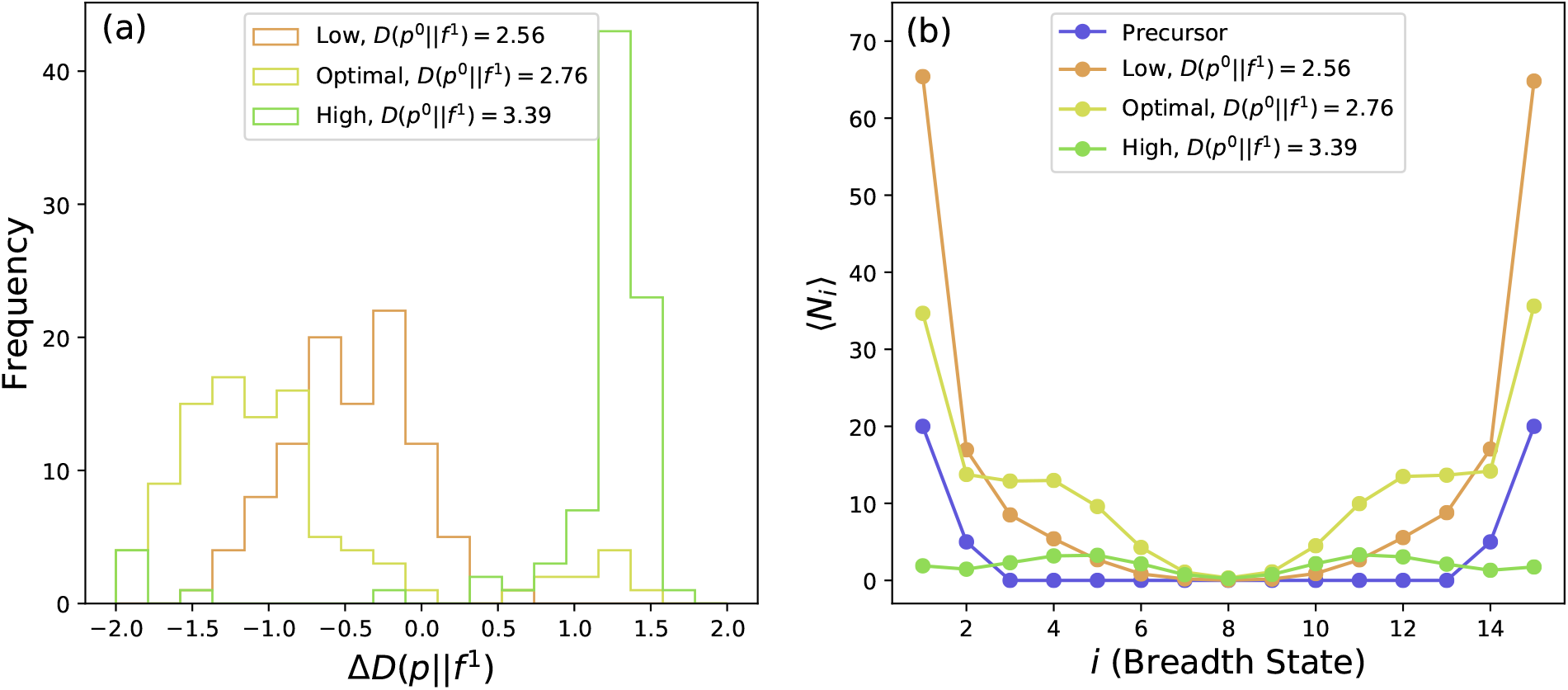
Optimal diversity of the B cell population. (a) For each GC, the change in relative entropy after prime immunization is recorded and graphed as a histogram for low (orange), optimal (yellow), and high (green) *D*(*p*^0^∥*f* ^1^). The GCs wherein the B cell population is extinguished experience a large increase of relative entropy (b) Decrease in relative entropy is a consequence of B cells on average populating higher breadth states after prime immunization.

Optimal *D*(*p*^0^∥*f* ^1^) (yellow, Fig. 6(a)) leads to larger decreases in the KLD within most of the GCs, indicating greater occupancy of higher breadth states in GCs (yellow, Fig. 6(b)) with respect to the precursor B cell population distribution (blue, Fig. 6(b)). A small number of GCs (*∼*10%) die and experience a KLD increase (yellow, Fig. 6(a)), indicating that those B cell populations have lost information about the antigenic environment. If *D*(*p*^0^∥*f* ^1^) (green, Fig. 6(a)) is too large, high rates of GC extinction lead to increases of the KLD within most of the GCs (*∼* 80%). The small number of GCs that survive do so because they manage to produce high breadth B cells by stochastic chance.

In order to understand the optimal choice of *f* ^2^ given *f* ^1^, for each B cell that exits as a bnAb after boost, we computed the breadth state of its ancestral B cell that initially seeded the GC at the beginning of boost. Fig. 7 shows the number of bnAb trajectories generated given that they originated from a starting sequence of breadth *i* after the prime. The optimal *D*(*p*^1^∥*f* ^2^) following immunization corresponding to lower than optimal *D*(*p*^0^∥*f* ^1^) must be large in order to generate bnAb trajectories from high breadth states (orange, Fig. 7) since occupancy within those states is very low after the sub-optimal prime (orange, Fig. 6(b)). For *D*(*p*^0^∥*f* ^1^) higher than the optimum, the optimal *D*(*p*^1^∥*f* ^2^) is small since the few GCs that survive prime mostly contain high breadth B cells (green, Fig. 6(b)). The best choice of *D*(*p*^1^∥*f* ^2^) following the optimal setting of *D*(*p*^0^∥*f* ^1^) falls between the optimal values of *D*(*p*^1^ *f* ^2^) for high and low values of *D*(*p*^0^∥*f* ^1^); i.e., *D*(*p*^1^∥*f* ^2^)_*high prime*_ *< D*(*p*^1^∥*f* ^2^)_*optimal prime*_ *< D*(*p*^1^∥*f* ^2^)_*low prime*_. The optimal setting for the boost successfully proliferates bnAb trajectories from several states near the highest breadth state (e.g. *i* = 5*—*7 or *i* = 9*—*11) that are populated by selection forces imposed by the optimal setting for the prime. That is, the optimal priming immunogen generates the right kind of B cell diversity, which makes possible many high probability evolutionary trajectories that mature into bnAbs. These trajectories ensue upon imposing the fitness landscape corresponding to the optimal boost.

**FIG 7:**
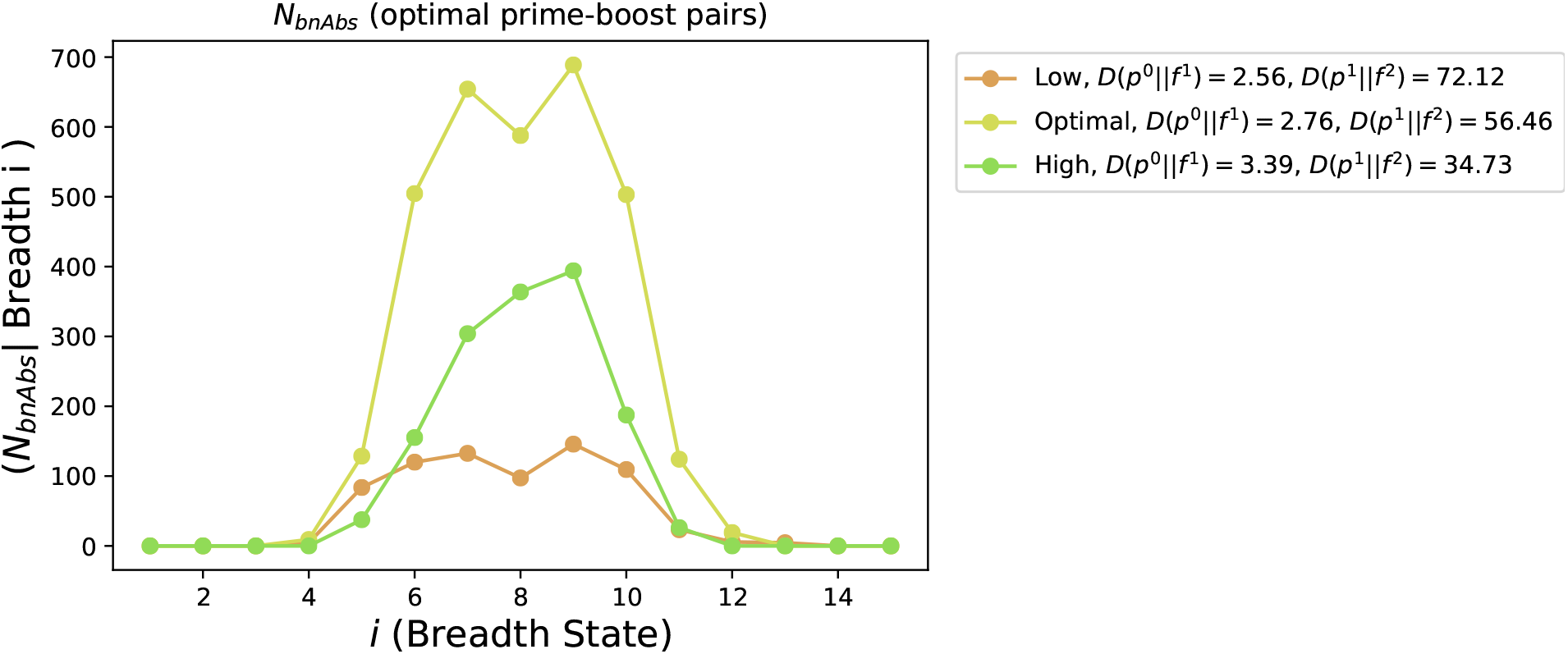
Characterizing optimal protocols. For the three prime protocols shown in Fig. 6, the numbers of evolutionary trajectories that exit the GCs as bnAbs after optimal boost are graphed as a function of the breadth state of the B cells that initially seed the trajectories at the beginning of boost.

### E. Phylogenetic analyses of evolving B cell populations

The phylogenetic trees shown in Figs. 8-10 tell a deeper story about the three possible immunization protocols analyzed in Figs. 6-7. For each of the 50 B cells that initially seed the GC, we compute the birth-death-mutation trajectories that emerge.

**FIG 8:**
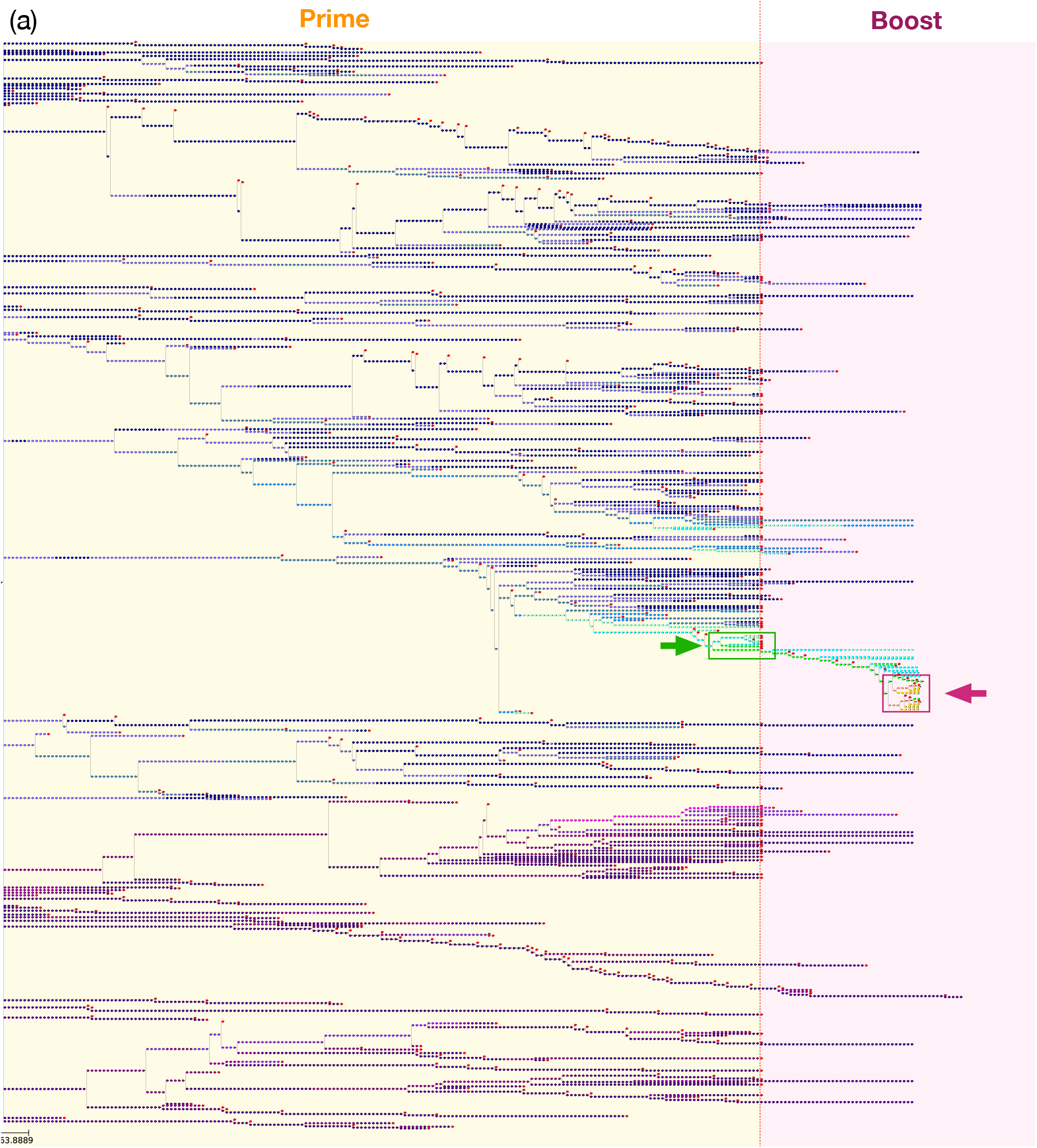
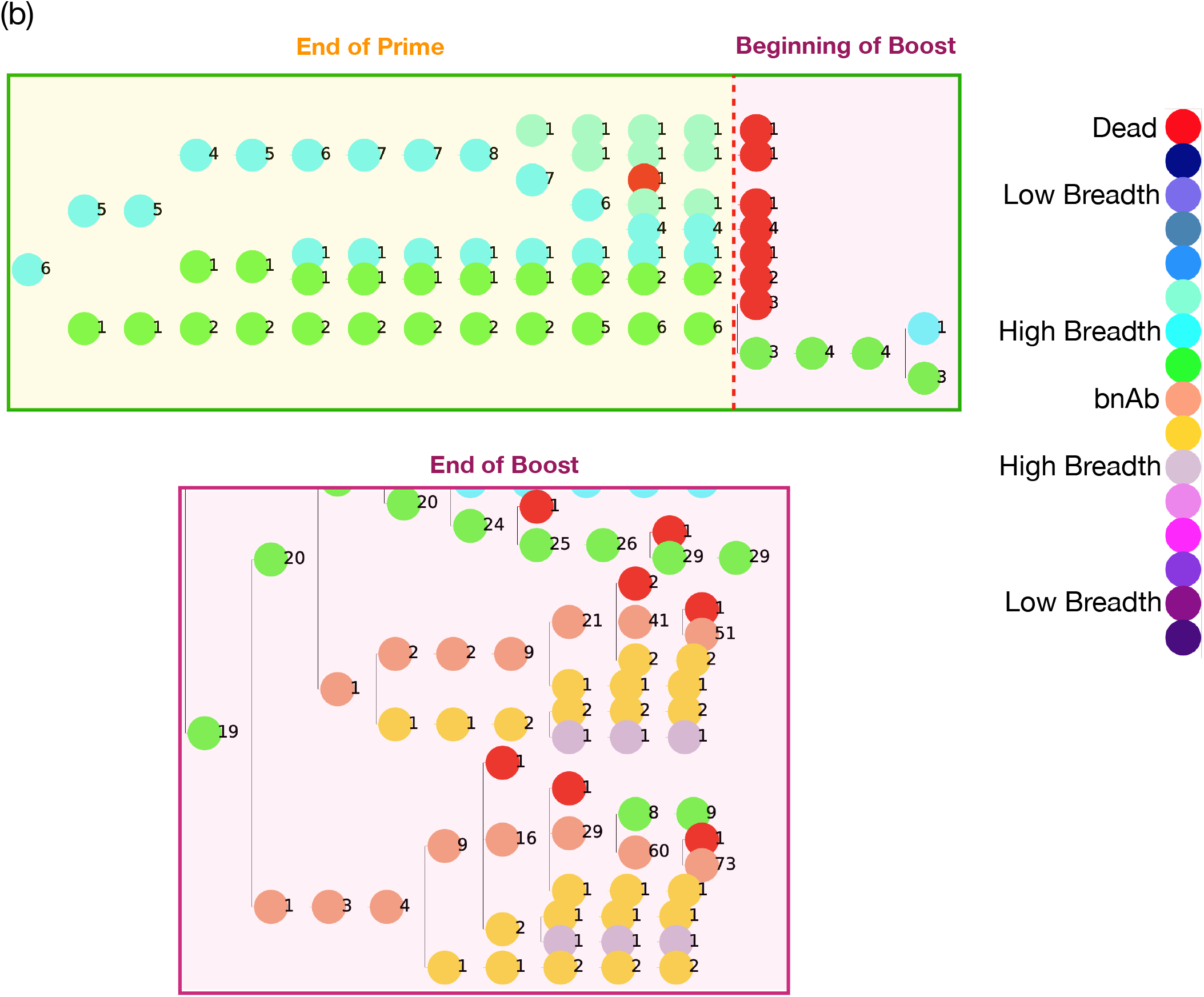
Phylogeny of evolving B cell lineages for low *D*(*p*^0^∥*f* ^1^) during prime followed by the optimal boost. (a) An example of a population of B cells evolving in a germinal center. The initial B cells were sampled from the precursor distribution (black, Fig. 2(b)) subjected to *D*(*p*^0^∥*f* ^1^) = 2.56 and *D*(*p*^1^∥*f* ^2^) = 72.12. The yellow region corresponds to the prime and the pink region to the boost. (b) The top panel is an expanded view of the green box (green arrow) in (a), which shows high breadth B cells generated at the end of prime and the beginning of boost. The bottom panel corresponds to the purple box (purple arrow) in (a), which occurs at the end of boost. Colors of circles depicting B cells denote their breadth states, and the numbers to the right of each circle quantify the number of B cells in that state.

**FIG 9:**
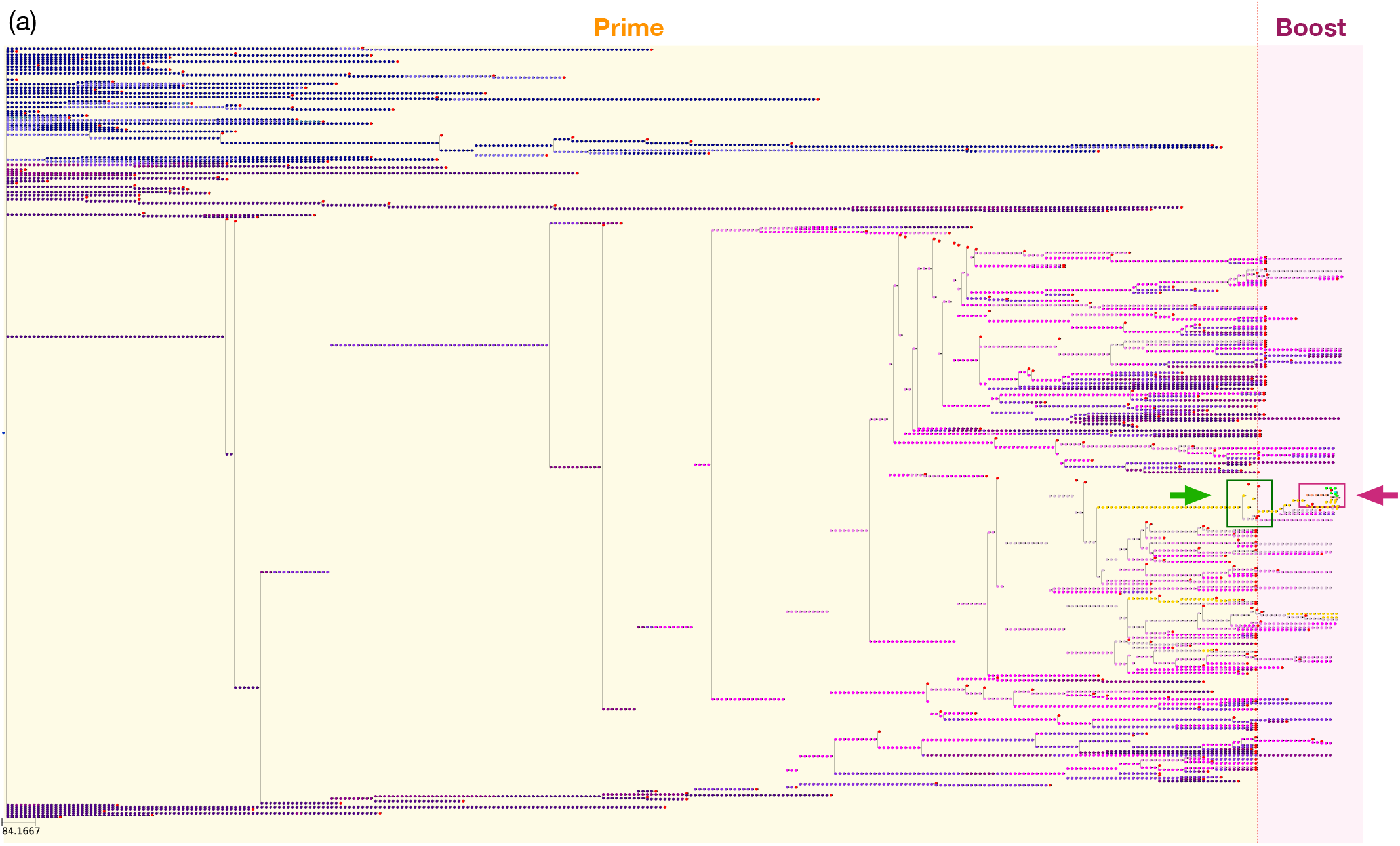
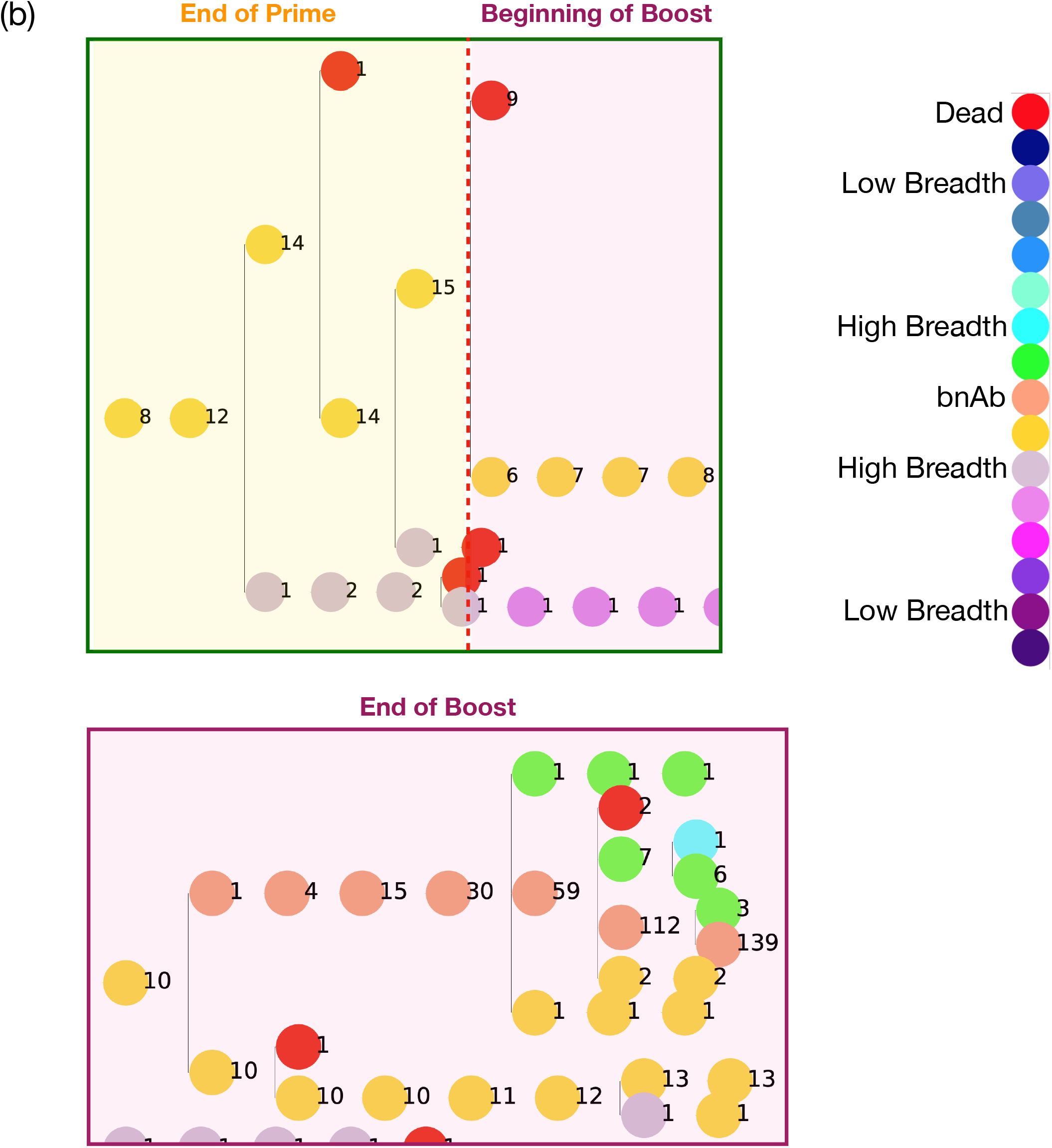
Phylogeny of evolving B cell lineages for high *D*(*p*^0^∥*f* ^1^) during prime followed by the optimal boost. (a) An example of a population of B cells evolving in a germinal center. The initial B cells were sampled from the precursor distribution (black, Fig. 2(b)) subjected to *D*(*p*^0^∥*f* ^1^) = 3.39 and *D*(*p*^1^∥*f* ^2^) = 34.73. The yellow region corresponds to the prime and the pink region to the boost. (b) The top panel is an expanded view of the green box (green arrow) in (a), which shows high breadth B cells generated at the end of prime. The bottom panel corresponds to the purple box (purple arrow) in (a), which corresponds to the end of boost. Colors of circles depicting B cells denote their breadth states, and the numbers to the right of each circle quantify the number of B cells in that state.

**FIG 10:**
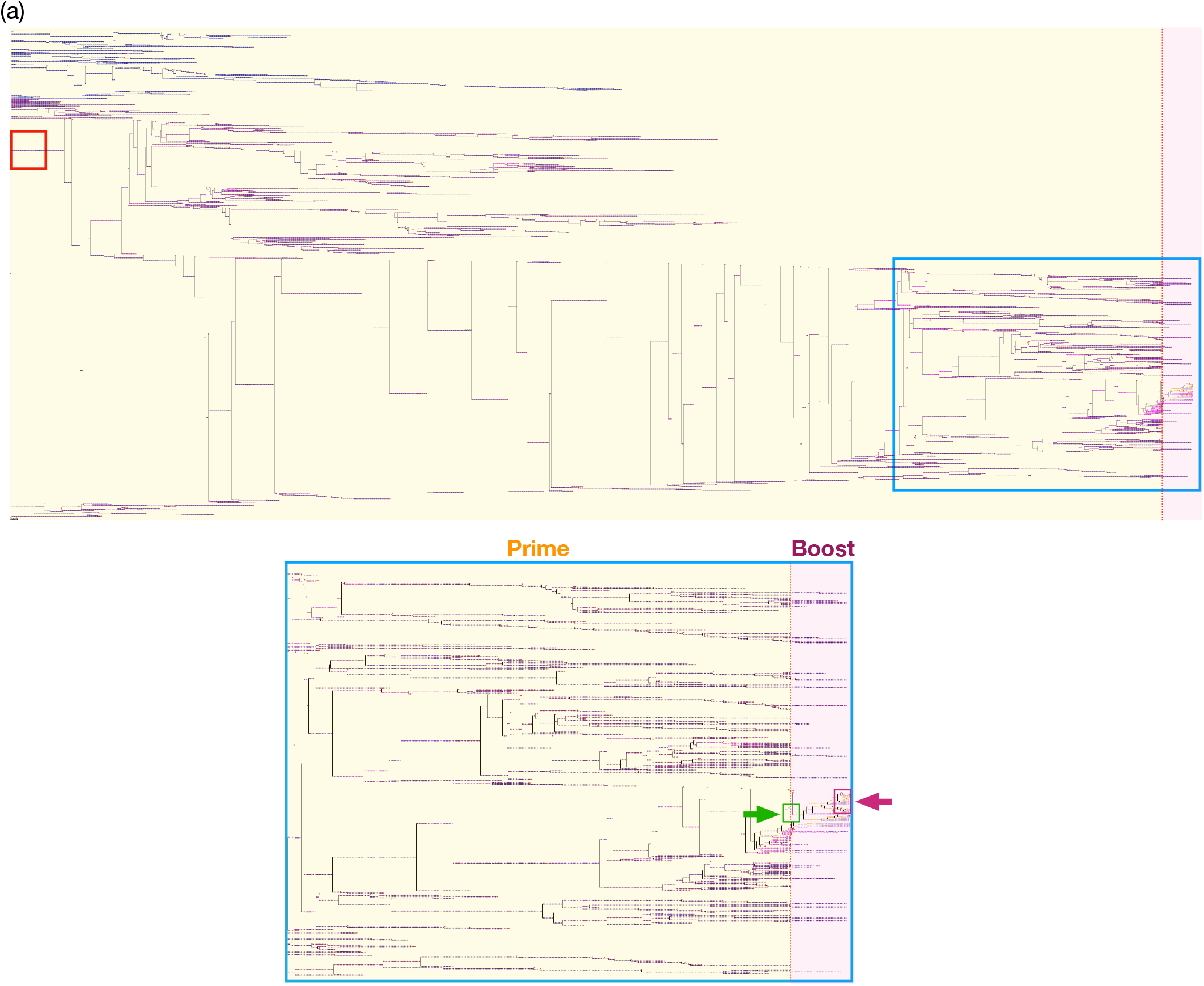
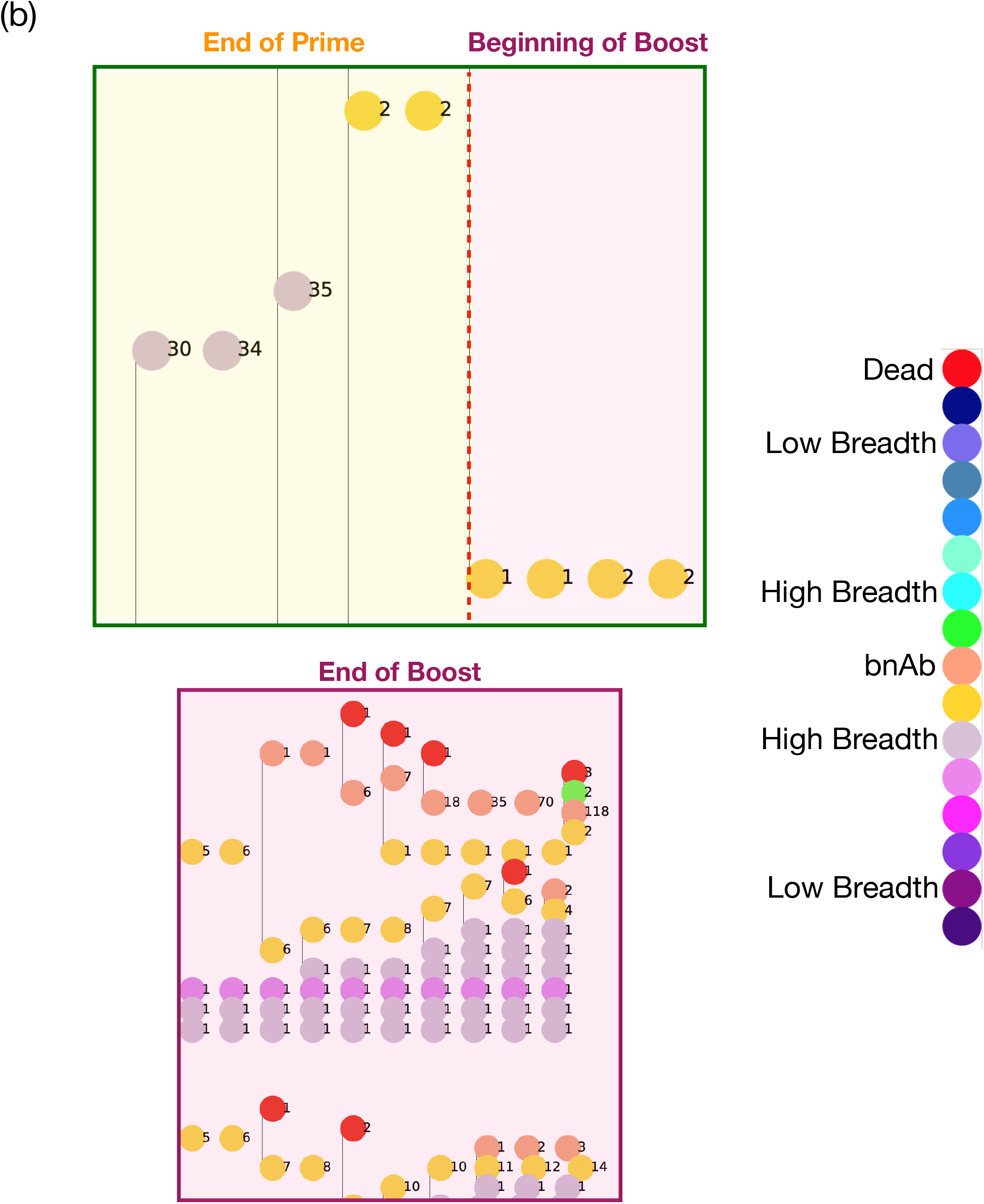
Phylogeny of evolving B cell lineages for optimal *D*(*p*^0^∥*f* ^1^) during prime followed by the optimal boost. (a) An example of a population of B cells evolving in a germinal center. The initial B cells were sampled from the precursor distribution (black, Fig. 2(b)) subjected to *D*(*p*^0^∥*f* ^1^) = 2.76 and *D*(*p*^1^∥*f* ^2^) = 56.46. The yellow region corresponds to the prime and the pink region to the boost. (b) The top panel is an expanded view of the green box (green arrow) in (a), which shows high breadth B cells generated at the end of prime. The bottom panel corresponds to the purple box (purple arrow) in (a), which corresponds to the end of boost. Colors of circles depicting B cells denote their breadth states, and the numbers to the right of each circle quantify the number of B cells in that state.

At low *D*(*p*^0^∥*f* ^1^), most of the initial crop of B cells manage to seed trajectories that survive until the end of prime (Fig. 8(a), yellow region). Yet, since selection is favorable for low breadth, these trajectories predominantly generate low breadth B cells; when a high *D*(*p*^1^∥*f* ^2^) is set during boost (see Fig. 8(a), pink region), nearly all of the trajectories sampled from these cells die. In rare cases, one of the precursor trajectories will stochastically manage to acquire enough breadth-enhancing mutations to generate high breadth cells before the end of prime. The top panel in Fig. 8(b) is an expanded view of the green box shown in Fig. 8(a). One of the trajectories leads to production of 6 high breadth B cells (green circles), which get selected at the beginning of boost and eventually lead to the proliferation of 124 bnAbs (orange circles, Fig. 8(b), bottom panel, an expanded view of the purple box in Fig. 8(a)). These rare events allow the GC to survive when selection for binding of conserved epitopes is low during prime and high through boost.

The opposite effect is observed at high *D*(*p*^0^||*f* ^1^). Under strong selection for high breadth, most trajectories generated by the precursor B cells die quickly during prime (see Fig. 9(a), yellow region). Yet, due to rare stochastic fluctuations, one of the precursor trajectories may generate a B cell of sufficiently high breadth that under strong selection generates a burst of replication and branching events (Fig. 9(a)); in the top panel of Fig. 9(b), these bursting events culminate in the production of 15 high breadth B cells (yellow circles) at the end of prime. During boost, these cells rapidly replicate and proliferate 139 bnAbs (orange circles, Fig. 9(b), bottom panel). Thus, low and high *D*(*p*^0^||*f* ^1^) protocols can induce bnAb production through rare events; the dynamics of these events differ depending on the strength of selection pressure during prime. The results are consistent with recent experiments [46] showing that homogeneous GCs are the product of clonal bursts (Fig. 9) though AM can take place in the presence or absence of such bursts (Fig. 8).

Phylogenetic trees for the optimal prime-boost protocol (Fig. 10(a)) present features which differ dramatically from the sub-optimal trees. Most noticeably, the tree exhibits considerably greater complexity with the presence of highly dense replication and mutation events. Due to the flux balance that prevents extinction and rapid proliferation of the population, GC lifetime during prime lasts significantly longer than for sub-optimal immunization. Thus, the precursor B cells follow longer trajectories that search breadth space more effectively. As a consequence, the probability of generating large numbers of higher breadth B cells is greatly enhanced. The top panel of Fig. 10(b) shows that, at the end of prime, a large number of high breadth sequences (*n ∼* 35, light purple circles) is produced. It is highly likely that at the beginning of boost (top panel, Fig. 10(b)), there is at least one B cell sufficiently close to the bnAb state (yellow circle) which eventually seeds the rapid proliferation of bnAbs by the end of boost (bottom panel, Fig. 10(b)).

An additional notable feature is evidence of a phenomenon akin to clonal interference. For the GC trajectory shown in Fig. 10, the 50 precursor B cells have the following distribution: *N*_1_ = 20, *N*_2_ = 4, *N*_14_ = 9, and *N*_15_ = 17. Interestingly, a single breadth 15 B cell (red box, Fig. 10(a)) manages to generate progeny that acquire breadth-enhancing mutations sooner than any of the other initially higher breadth cells. As a result, significant clonal expansion of this B cell lineage outcompetes trajectories produced by all other precursor cells. Eventually, all of the B cells that exit prime and seed the GC during boost originate from this single precursor cell (blue box, Fig. 10(a)). The exhaustive search process during optimal prime immunization leads to occupancy of high breadth states that maximize the likelihood of proliferating trajectories which eventually find the bnAb state during boost (bottom panel, Fig. 10(b)).

## III. CONCLUSIONS

Developing effective vaccines and immunization protocols to induce the production of antibodies that neutralize diverse strains of highly mutable viruses is a major global health challenge. To date, a vaccination strategy that can generate such bnAbs against viruses like HIV or influenza have not been developed. Antibodies are produced by the Darwinian evolutionary process of AM. The immunogens used in a vaccination strategy impose selection forces on the B cell population. Generating bnAbs requires controlling selection forces so that generalists which can neutralize diverse mutant strains of the virus, rather than specialists that neutralize specific strains, evolve. In this paper, we combined stochastic simulations of a minimal model for AM with an information theoretic metric to understand the mechanistic reasons underlying why certain vaccination protocols are optimal for generating bnAbs.

We studied a case where two immunizations were allowed, a prime and boost. Thus, immunization protocols must drive the evolution of bnAbs in a non-adiabatic way. We first defined a quantitative measure of the thermodynamic selection force that acts on the existing B cell population upon immunization, and which thus drives the non-equilibrium fluxes that affect the evolution of the population [15, 17]. The KLD, *D*(*p*^*j*^||*f*^*j*+1^), measures the strength of this force. Alternatively, from the standpoint of statistical learning theory, the KLD can be interpreted as a cost function that the B cell population attempts to minimize by replicating and mutating to higher breadth states. Therefore, the KLD also sets the learning rate per injection during training. If it is set too low, optimization is slow and minimal learning happens before all of the antigen is consumed. If it is too high, unexpected divergent behavior emerges and the KLD rapidly increases before extinction.

We find that there is an optimal combination of KLD’s imposed by prime and boost that maximizes the generation of bnAbs. Optimal prime occurs when the fitness of the lowest breadth states falls below the basal death rate, leading to a balance of fluxes that causes neither extinction nor rapid proliferation of the population. This provides the B cells a sufficient amount of time to acquire the low-probability breadth enhancing mutations necessary to optimally increase diversity and maintain high survival probability. For a given KLD that defines the prime, the optimal KLD for the boost is always higher. These results are consistent with previous studies of the effects of changing prime-boost immunization strategies by altering mutational distances, concentrations and other features of the immunogens [2, 16, 17, 27].

Importantly, our study provides several new insights in terms of the character of the interesting non-equilibrium process under consideration. If the KLD upon priming or boosting is not too large, the B cell population evolves in time to try to adapt to the imposed selection force, resulting in reduction of the KLD with time. However, perfect adaptation to the selection forces cannot occur because of the finite amount of antigen supplied by the injection. Thus, unless there is extinction of the B cell population or continuous provision of antigen, the distribution eventually relaxes to a non-equilibrium steady-state where, as expected, the thermodynamic selection force, *D*(*p*(*t*) ||*f*), approaches a positive and constant value. Non-equilibrium thermodynamics tells us that if the force is larger, adaptation should occur faster because of a larger irreversible flux [47]. Contrary to this expectation, if the initially imposed KLD upon the prime or boost immunization is too large, the B cell population is extinguished with high probability. Thus, no adaptation is possible if the non-equilibrium driving force is too large.

If the KL divergence is too small during the prime, most B cell lineages successfully survive, but they do not gain much information about the bnAb state. These lineages mostly die during the boost, which has to be very aggressive in order for any bnAbs to evolve. The few bnAbs that do evolve are the product of rare stochastic trajectories that acquire the right breadth-enhancing mutations to become nearly bnAbs during the prime. These are then rapidly expanded during the boost. If the KL divergence during the prime is too large, most B cell lineages become extinct because their fitness is very low. Again, the B cell population does not gain information about the bnAb state. The few bnAbs that ultimately evolve are again the product of rare fortunate trajectories that survive strong selection pressure during prime and access the bnAb state during the boost. In the latter case, the dynamics are ballistic-like as shown by the phylogenetic trees (Fig. 9(a)). The optimal priming condition sets the B cell population off-equilibrium sufficiently so that B cells must evolve to higher breadth states to proliferate effectively, but the population does not become extinct with high probability. This balanced selection force during the prime results in complex evolutionary trajectories that generate B cells in states close to the bnAb state. During the boost, these lineages can evolve into bnAbs. In the process, there is evidence of clonal interference. Therefore, the optimal prime results in the B cell population acquiring information about the bnAb state and the right kind of diversity of lineages is generated.

Recently, it has been shown that phylogenetic trees exhibit different topological asymptotic scaling laws which depend on whether they are balanced or unbalanced [48]. Upon visual inspection of Figs. 8-10, it is clear that the trees produced by the optimal protocol exhibit a vastly different topology from those constructed by sub-optimal protocols. A scaling analysis could reveal that optimal and sub-optimal protocols lead to different asymptotic behaviors with respect to phylogeny, a result that could reflect a complex interplay between ecological and evolutionary processes during AM [49].

We hope our results will guide more detailed simulations and experiments designed to generate immunogens that can impose an optimal prime/boost protocol. We also hope that our work will motivate theoretical studies on how evolutionary forces can select for generalists in realistic conditions that are not periodic oscillations in environments.

