## Supplementary Material for "Mechanisms underlying vaccination protocols that may optimally elicit broadly neutralizing antibodies against highly mutable pathogens"

(Dated: February 2, 2021)

### I. GILLESPIE REACTIONS

The ordinary differential equations described by Eq. (2) in the main text were converted into the following set of "chemical reactions" and solved using the Gillespie method [1, 2]:

For the edge states $i = 1$ and $i = 15$,

$\mathrm{N}_1 \xrightarrow{\mathrm{f}_1} 2\,\mathrm{N}_1$

$\mathrm{N}_1 \xrightarrow{\mu_{1,2}} \mathrm{N}_2$

$\mathrm{N}_1 \xrightarrow{\mu_{1,0}} \mathrm{N}_0$

$\mathrm{N}_{15} \xrightarrow{\mathrm{f}_{15}} 2\,\mathrm{N}_{15}$

$\mathrm{N}_{15} \xrightarrow{\mu_{15,14}} \mathrm{N}_{14}$

$\mathrm{N}_{15} \xrightarrow{\mu_{15,0}} \mathrm{N}_0$

For all states between $i = 1$ and $i = 15$,

$\mathrm{N}_i \xrightarrow{\mathrm{f}_i} 2\,\mathrm{N}_i$

$\mathrm{N}_i \xrightarrow{\mu_{i,i-1}} \mathrm{N}_{i-1}$

$\mathrm{N}_i \xrightarrow{\mu_{i,i+1}} \mathrm{N}_{i+1}$

$\mathrm{N}_i \xrightarrow{\mu_{i,0}} \mathrm{N}_0$

Stochastic trajectories stop when only species $N_0$ remains ($\sum_{i\neq 0} N_i(t) = 0$) or when the maximum population size is reached e.g. $\sum_{i\neq 0} N_i(t) = 200$ for the calculations shown in the main text.

### II. EFFECTS OF INCREASING DISCRETIZATION

Since exploring the continuum limit is analytically intractable, additional calculations with a larger number of bins were carried out. As germ-line B cells typically have low breadth, precursor B cells are sampled from a log-normal distribution with $\mu = 3.0$ and $\sigma = 1.0$ (Fig. S1). The simulation protocol described in the main text is carried out with the following parameters: $K = 32$, $\mu_{i0} = 0.01$, and $\mu_{ij} = 0.05$. The mutation rates $\mu_{i,i+1} = 0.3\mu_{ij}$ and $\mu_{i+1,i} = 0.7\mu_{ij}$ if $i < K/2$. Conditions flip for $i > K/2$ and for $i = K/2$,

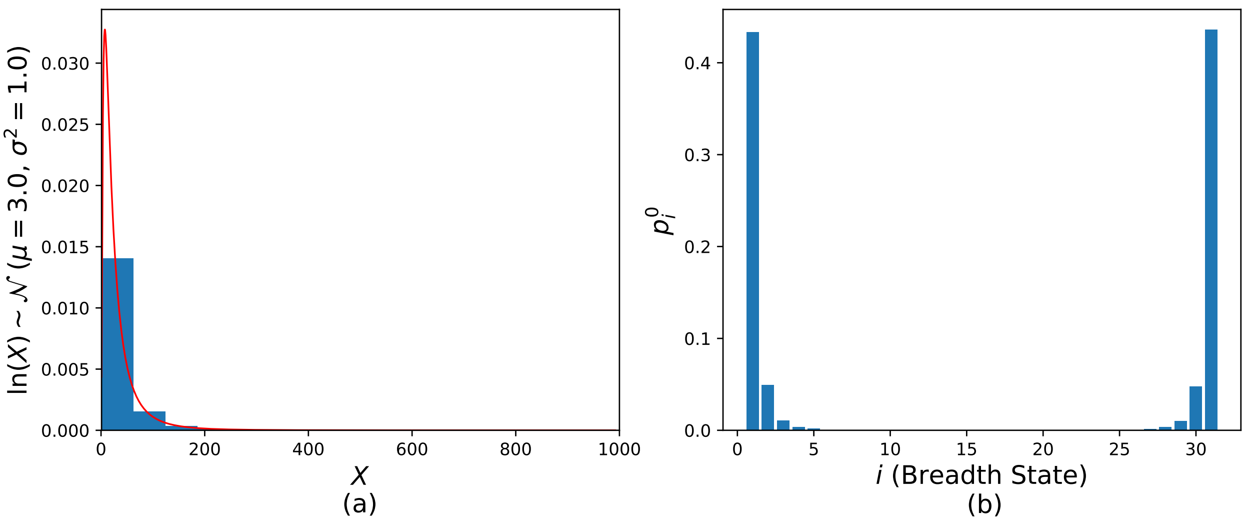


FIG. S1: Initial distribution of B cells. (a) Precursor B cells are sampled from a log-normal distribution with $\mu = 3.0$ and $\sigma = 1.0$. (b) Normalizing with respect to K = 31 bins yields the precursor B cell distribution $p_i^0$.

$\mu_{i,i+1} = \mu_{i,i-1} = 0.5\mu_{ij}$. As the number of bins increases, the overall fitness per bin decreases and the other parameters must change accordingly. Yet, the same assumptions described in the main text regarding the parameter regime hold.

Fig. S2 shows the number of bnAbs produced per GC for three points near the optimal setting of $D(p^0||f^1)$ and all possible subsequent settings of $D(p^1||f^2)$. Since the number of bins has increased, we observe the titers in bins $14 - 18$, the states of highest possible breadth. We clearly observe that an optimal protocol exists and $D(p^1||f^2) > D(p^0||f^1)$ at the optimal points of all protocols. As was the case with $K = 16$ bins, Fig. S3 shows that optimal prime immunization occurs at the point just after the onset of GC collapse (Fig. S3(a)) when fitness in the lowest breadth states drops below the death rate (Fig. S3(b)).

### III. ADIABATIC LIMIT AND EXTINCTION

Fig. S4 shows the adiabatic limit of the vaccination protocols presented in Fig. 5 of the main text. Fig. S4(a) shows a small sample of GC trajectories when $D(p^0||f^1)$ is near the optimal point; Fig. S4(b) shows the rare surviving trajectories when $D(p^0||f^1)$ is too high. If antigen is continuously pumped into the germinal center, there is no constraint on the maximum size of the B cell population. As the population increases in size, the

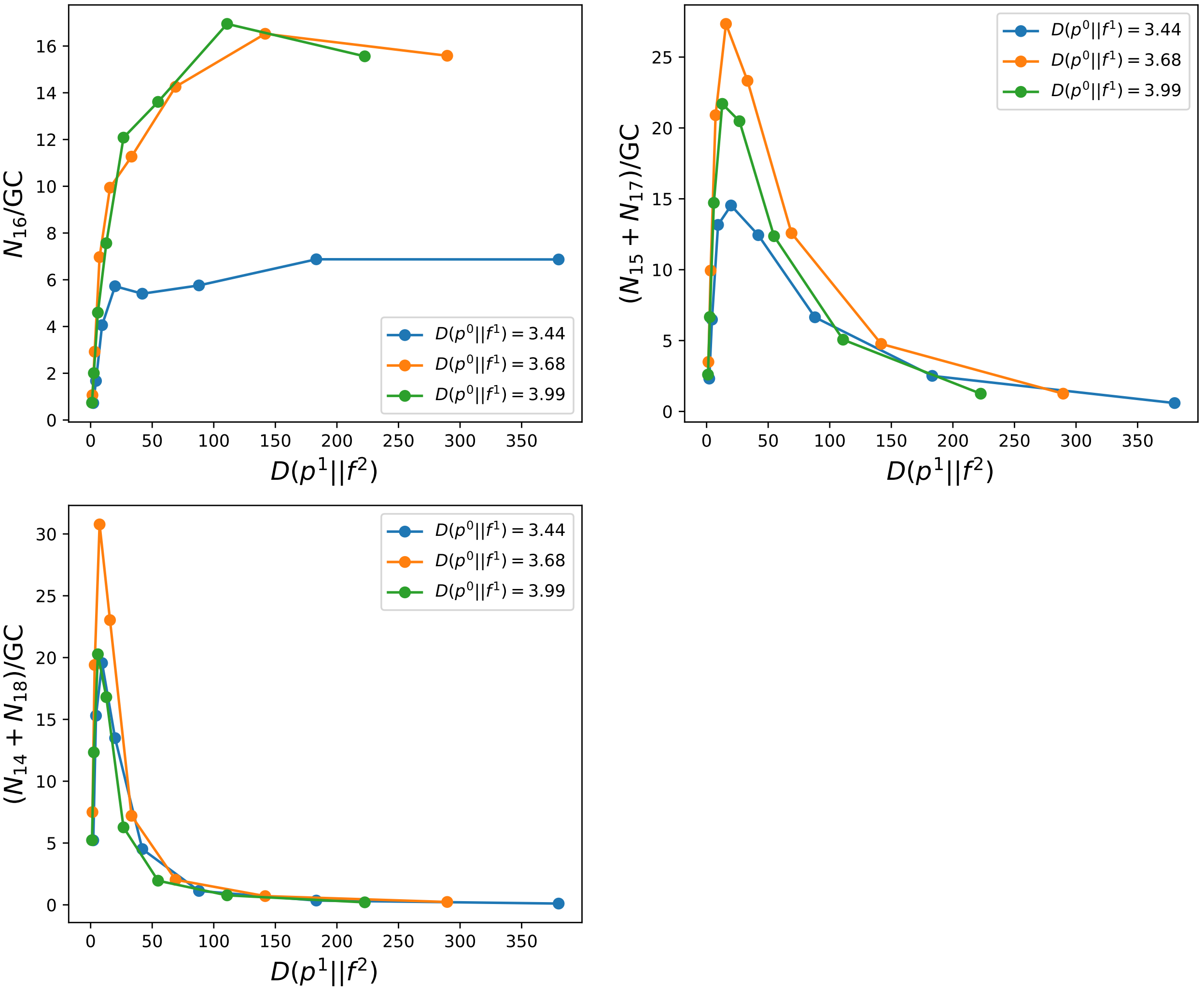


FIG. S2: Response to different vaccination protocols. For each prime immunization choice of KL divergence, $D(p^0||f^1)$ (colored curves), the number of high breadth B cells produced per GC is shown over a range of boost immunization choices, $D(p^1||f^2)$.

KLD decreases (Fig. S4) until it relaxes to a near equilibrium state ($D(p(t)||f^1) \sim 0$). The relaxation time tends to decrease as the selection force $D(p^0||f^1)$ increases.

Fig. S5 shows the behavior of $D(p(t)||f^1)$ within GC trajectories that result in extinction events. Along these trajectories, strong selection forces induce flux to low breadth states and the death state. As a consequence, the population distribution evolves to be further away from the imposed fitness and $D(p(t)||f^1)$ increases. Near the optimal point (Fig. S5(a)), there is a prolonged time until extinction since the learning rate $D(p^0||f^1)$ is only slightly larger than the optimal choice. As a result, B cells in most of the GCs manage to successfully reduce the KLD and learn about the antigenic environment (Fig. S4(a)). If the learning

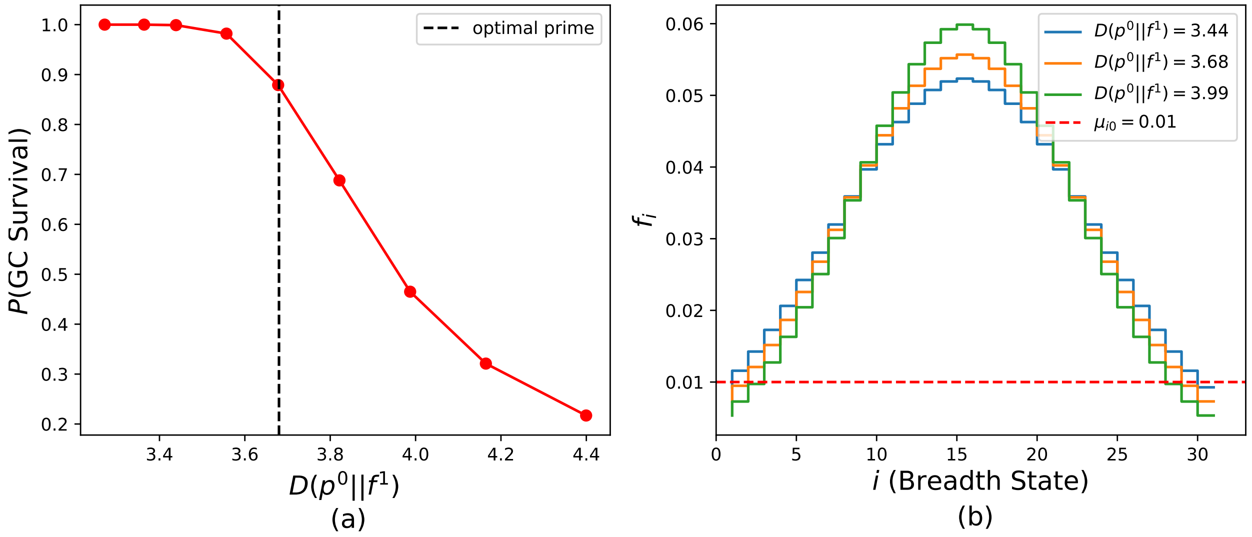


FIG. S3: Germinal center (GC) survival (B cell population is not extinguished). (a) Fraction of GCs that survive prime immunization for a range of values of $D(p^0||f^1)$. The fraction falls to $\sim 0.9$ at the optimal setting and drops significantly beyond this point. (b) Sample fitness landscapes at the points located on the curve shown in (a).

rate is too high (Fig. S5(b)), we observe rapid divergent behavior in the KLD.

[1] D. T. Gillespie, A general method for numerically simulating the stochastic time evolution of coupled chemical reactions, Journal of computational physics **22**, 403 (1976).

[2] D. T. Gillespie, Exact stochastic simulation of coupled chemical reactions, The journal of physical chemistry **81**, 2340 (1977).

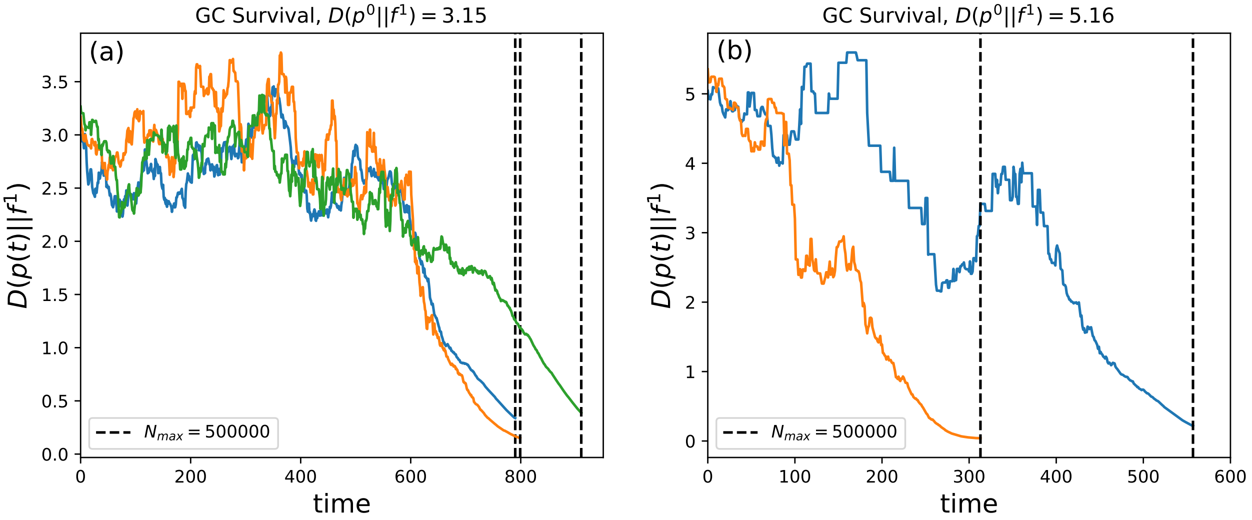


FIG. S4: $D(p(t)||f^1)$ of surviving GC trajectories when (a) $D(p^0||f^1) = 3.15$ and (b) $D(p^0||f^1) = 5.16$. To explore the adiabatic limit, the maximum population size is increased to $N_{max} = 500000$.

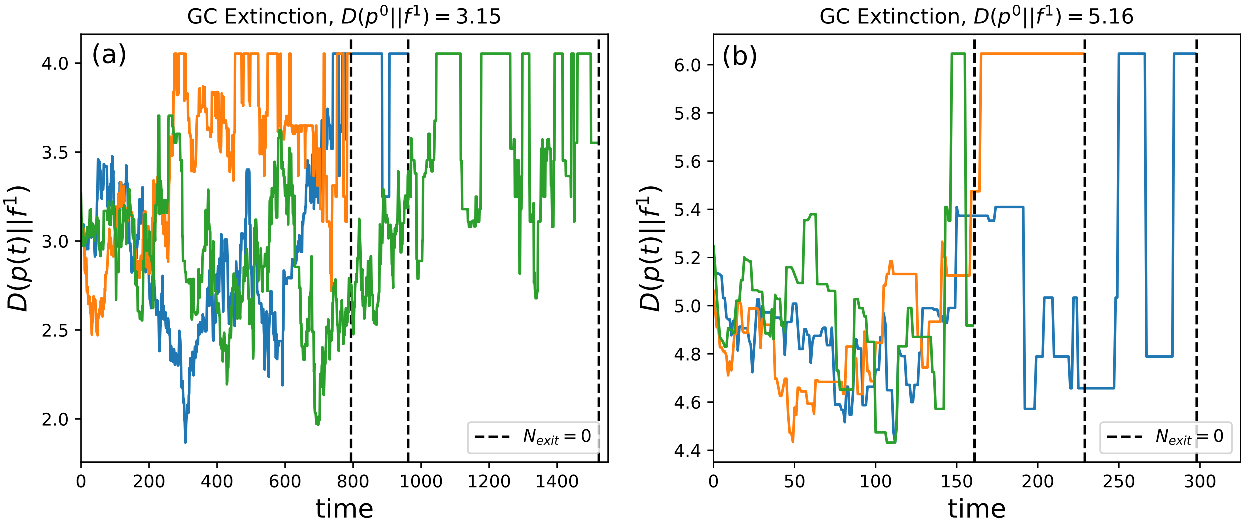


FIG. S5: $D(p(t)||f^1)$ within GC trajectories that lead to population extinction when (a) $D(p^0||f^1) = 3.15$ and (b) $D(p^0||f^1) = 5.16$.
